# CDCA7 facilitates MET1-mediated CG DNA methylation maintenance in centromeric heterochromatin via histone H1

**DOI:** 10.1101/2025.09.22.677529

**Authors:** Shuya Wang, Tong Li, Matthew Naish, Russell Chuang, Evan K. Lin, Christian Fonkalsrud, He Yan, Suhua Feng, Ian R. Henderson, Steven E. Jacobsen

**Affiliations:** Molecular Biology Institute, University of California Los Angeles, Los Angeles, CA 90095, USA; Department of Molecular, Cell and Developmental Biology, University of California Los Angeles, Los Angeles, CA 90095, USA; Department of Plant Sciences, University of Cambridge, Cambridge, CB2 3EA, UK; Eli & Edythe Broad Center of Regenerative Medicine & Stem Cell Research, University of California at Los Angeles, Los Angeles, CA 90095, USA; Howard Hughes Medical Institute, University of California Los Angeles, Los Angeles, CA 90095, USA

## Abstract

DNA methylation is a conserved epigenetic modification essential for maintaining genome stability. However, how methyltransferases maintain CG methylation within compact chromatin, including centromeres, remains unclear. In humans, CDCA7 is necessary for the inheritance of DNA methylation at juxta-centromeres. Mutations that impair its ability to bind chromatin result in Immunodeficiency, Centromeric Instability, and Facial Anomalies (ICF) syndrome, characterized by centromeric instability. To investigate whether CDCA7 function is conserved, we identified two *Arabidopsis thaliana* orthologs, *CDCA7A* and *CDCA7B*. The loss of both copies results in CG hypomethylation at pericentromeric regions and centromeric satellite repeat arrays. Machine learning analysis suggested that heterochromatic nucleosomes, with enrichment of H1, H2A.W, and H3K9me2 levels, depend heavily on CDCA7 proteins for CG methylation maintenance of the associated DNA. Loss of H1 restores heterochromatic DNA methylation in *cdca7a cdca7b* mutants, indicating that *CDCA7A* and *CDCA7B* mainly remodel H1-containing nucleosomes for methyltransferases to access DNA. Notably, in *h1.1 h1.2* mutants, CG methylation shows a significant increase in centromeres, which reveals a new inhibitory role of H1 in DNA methylation maintenance within satellite repeat arrays. Centromeric DNA hypermethylation is lost in *h1.1 h1.2 cdca7a cdca7b* quadruple mutants, demonstrating that *CDCA7A* and *CDCA7B* can act independently of H1 to enhance MET1 activity. Overall, these findings establish *CDCA7A* and *CDCA7B* as conserved regulators of DNA methylation within heterochromatin and centromeric satellite repeat arrays.

## Introduction

DNA methylation plays an essential role in silencing genes and transposable elements (TEs), which is indispensable for development and reproduction in mammals [1-3]. In *Arabidopsis thaliana* (hereafter *Arabidopsis*), DNA methylation occurs in CG, CHG (H = A, T, or C), and CHH sequence contexts. VIM proteins recognize hemi-methylated cytosines and recruit MET1 to maintain CG methylation, while non-CG methylation is propagated by CHROMOMETHYLASE2 (CMT2) and CHROMOMETHYLASE3 (CMT3) [4]. The RNA-directed DNA methylation (RdDM) pathway *de novo* methylates TEs via methyltransferases DRM1 and DRM2 [5]. DNA methylation maintenance faces a significant challenge in regions with a high density of nucleosomes, particularly in heterochromatin, which restricts access to un- or hemi-methylated cytosine substrates for DNA methyltransferases [6-9]. To overcome this barrier, *Arabidopsis* DDM1 has been proposed to remodel H1-containing nucleosomes, enabling DNA methyltransferases to access pericentromeric chromatin [6, 7, 10-12]. However, the mechanism guiding DDM1 to its nucleosomal targets remains unclear.

In mammals, HELLS (the homolog of DDM1) depends on CDCA7 for localization and activation [13]. CDCA7 contains an evolutionarily conserved zf-4CXXC_R1 domain that recognizes hemi-methylated CpG in the DNA major groove [13-17]. Simultaneously, CDCA7 can interact with HELLS and relieve its catalytic autoinhibition [13]. Current research suggests that CDCA7 recruits HELLS to satellite DNA arrays via the zf-4CXXC_R1 domain and activates HELLS to remodel nucleosomes for replication-uncoupled DNA methylation maintenance via UHRF (an ortholog of VIM) and DNMT1 (an ortholog of MET1) [15]. Mutations in the zf-4CXXC_R1 domain, or HELLS-interacting domain, of CDCA7 lead to DNA hypomethylation at juxta-centromeric satellite DNA, resulting in centromere instability and ICF syndrome [18]. Mammalian centromeres consist of megabase arrays of alpha-satellite repeats, which serve as the site for CENP-A histone loading and kinetochore formation [19-22]. Although the *Arabidopsis* genome is smaller (∼130 Mb) compared to the human genome (∼3 Gb), it also uses comparably sized megabase arrays of a 178-base pair satellite repeat (*CEN178*) for its centromeres [23-25]. Centromeric and pericentromeric regions are heavily methylated at CG sites in both mammals and *Arabidopsis*. However, they differ in that CENP-A-occupied repeats are CG hypomethylated in mammals, whereas *Arabidopsis* CENH3-occupied *CEN178* repeats are densely CG methylated [22, 23]. Whether CDCA7 proteins operate through similar mechanisms to assist CG methylation maintenance in plant centromeres and other genomic regions remains unknown. Additionally, DDM1 can remodel nucleosomes independently, while HELLS requires CDCA7 binding to perform remodeling activities [11, 26]. This indicates that *Arabidopsis* CDCA7 proteins may function differently from their mammalian counterparts during DNA methylation maintenance.

In this study, we show that two Class I CDCA7 proteins work redundantly to maintain CG methylation in *Arabidopsis* centromeric satellite repeat arrays and pericentromeric regions. The DNA CG hypomethylation seen in *cdca7a cdca7b* mutants is less severe than in the *ddm1-2* and *met1* mutants, suggesting that CDCA7 partially contributes to DDM1-dependent methylation. Machine learning analysis identified nucleosome density, histone H1 enrichment, H2A.W abundance, and H3K9me2 levels as key chromatin features predicting methylation loss in the *cdca7a cdca7b* background, indicating that compact chromatin depends heavily on CDCA7 for CG methylation maintenance. Supporting this, histone H1 depletion restores DNA methylation in *cdca7a cdca7b* mutants, showing that H1 is the main barrier to VIM and MET1 access when CDCA7 proteins are absent. Therefore, in the wild-type, *CDCA7A* and *CDCA7B* act on H1-containing nucleosomes to promote the access of VIM and MET1. To determine whether CDCA7 functions independently of H1, we examined the *h1.1 h1.2* mutant. We found significant centromeric CG hypermethylation, revealing an unexpected role for H1 in preventing CG methylation within the *Arabidopsis* centromere satellite repeat arrays. This DNA hypermethylation is abolished in *h1.1 h1.2 cdca7a cdca7b* quadruple mutants, indicating that CDCA7 also promotes CG methylation, even in the absence of H1. These findings highlight the conserved role of CDCA7 orthologs and demonstrate that *Arabidopsis* CDCA7 supports MET1 activity across a range of nucleosome contexts, including centromere satellite repeat arrays. Our results also provide genetic evidence that histone H1 regulates DNA methylation levels in centromeric satellite repeat arrays.

## Results

### *CDCA7A* and *CDCA7B* are conserved CDCA7 orthologs

Mutations in three critical amino acids (R274, G294, and R304) of the CDCA7 zf-CXXC_R1 domain reduce its binding affinity to hemi-methylated CpG sites, leading to ICF syndrome [15]. Among the three classes of *Arabidopsis* CDCA7 proteins (Fig. 1A), only Class I CDCA7s (*CDCA7A* and *CDCA7B*) retain these residues and the cysteines required for zf-CXXC_R1 domain folding (Fig. 1B-C) [14]. Consistent with this conservation, *CDCA7A* and *CDCA7B* preferentially bind hemi-methylated DNA over fully methylated or unmethylated DNA *in vitro*, suggesting a similar selectivity *in vivo* [15]

**Figure 1.**
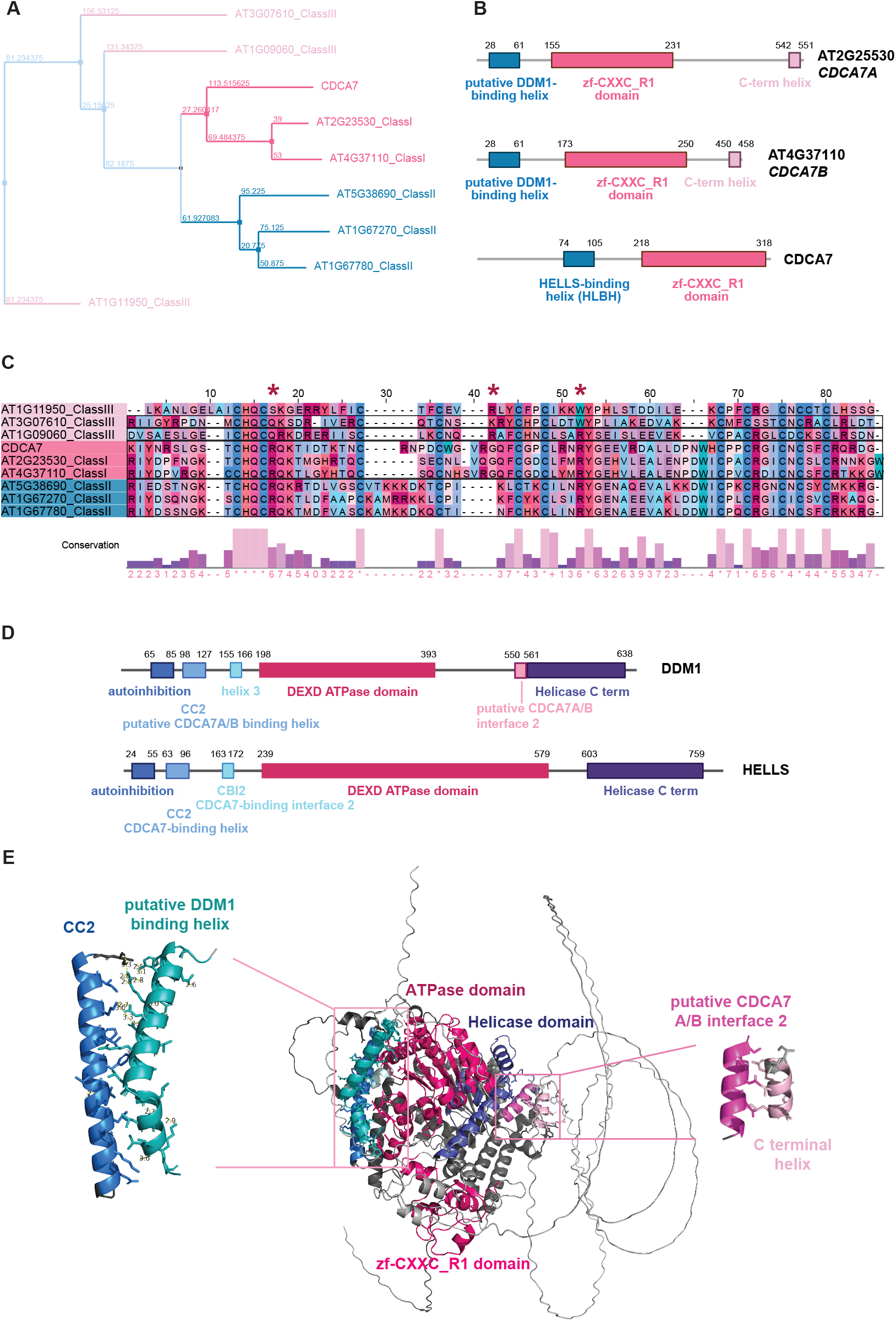
*CDCA7A* and *CDCA7B* are conserved CDCA7 orthologs. **A**. Phylogenetic tree based on the conservation of zf-CXXC_R1 domains of *homo sapiens* CDCA7 and its *Arabidopsis thaliana* homologs. **B**. Functional domain annotations of *homo sapiens* CDCA7 and two *Arabidopsis* Class I CDCA7s. **C**. Sequence alignment calculated by Clustal Omega of the zf-CXXC_R1 domains of *homo sapiens* CDCA7 and its *Arabidopsis* homologs. The red asterisks indicate key residues (R274, G294, and R304) that were mutated in ICF syndrome. **D**. Domain annotations of HELLS and DDM1. **E**. *CDCA7A* and DDM1 interaction model predicted by AlphaFold 3. Cyan represents the putative DDM1-binding helix of *CDCA7A*. Blue represents CC2 of DDM1. Light pink represents the C-terminal helix of *CDCA7A*. Pink represents putative CDCA7 interface 2 of DDM1. Dark pink represents the ATPase domain of DDM1. Purple indicates the Helicase C term domain of DDM1. Bright pink represents the zf-CXXC_R1 domain of *CDCA7A*.

Class I CDCA7s also share a similar protein structure with mammalian CDCA7, including an N-terminal helix analogous to the HELLS-binding helix (HLBH) (Fig. 1B, S1A). Both HELLS and *Arabidopsis* DDM1 have an N-terminal coiled-coil domain (CC2), which is known in HELLS to be critical for interaction with CDCA7 (Fig. 1D) [11, 15]. AlphaFold 3 (AF3) predicts that the HLBH of *CDCA7A* and *CDCA7B* directly interacts with DDM1’s CC2 through multiple types of interactions (Fig. 1E, S1B-D, Table S1-2) [27]. Consistent with this, DDM1 lacking the CC2-containing N-terminal domain cannot complement *ddm1* mutants [12], indicating that interaction with CDCA7A/B is essential for DDM1 function. In addition to the HLBH-CC2 interaction, HELLS also relies on the CDCA7-binding interface 2 (CBI2) to strengthen its association with CDCA7. By contrast, DDM1 is predicted to use its helicase domain to interact with the C-terminal helix of *CDCA7A/B* (Fig. 1E). This interaction likely compensates for the absence of CBI2, suggesting different recruitment strategies of CDCA7 proteins in plants and mammals. Overall, AF3’s structural predictions suggest that *CDCA7A* and *CDCA7B* retain key features of human CDCA7 and imply their functional conservation.

### *CDCA7A* and *CDCA7B* are required for the maintenance of heterochromatic DNA methylation

Given the evolutionary conservation between *Arabidopsis CDCA7A* and *CDCA7B* homologs and mammalian CDCA7, we investigated their roles in DNA methylation maintenance. Using CRISPR-Cas9, we generated loss-of-function mutants for *cdca7a, cdca7b, cdca7a cdca7b+/-*, and *cdca7a cdca7b* in the Col-0 background (Fig. S2A) and conducted Whole-Genome Bisulfite Sequencing (WGBS) to assess genome-wide DNA methylation. Loss of *CDCA7A* alone results in a 2% reduction in overall CG methylation, primarily observed in heterochromatin (Fig. 2A–E). In comparison, knocking out *CDCA7B* alone causes a 6% decrease in CG methylation at pericentromeric regions (Fig. 2A-E), while non-CG methylation shows less than 1% difference (Fig. 2A, Fig. S2B-C). These findings suggest that *CDCA7B*, compared to *CDCA7A*, plays a more prominent role in supporting VIM and MET1-mediated CG methylation.

**Figure 2.**
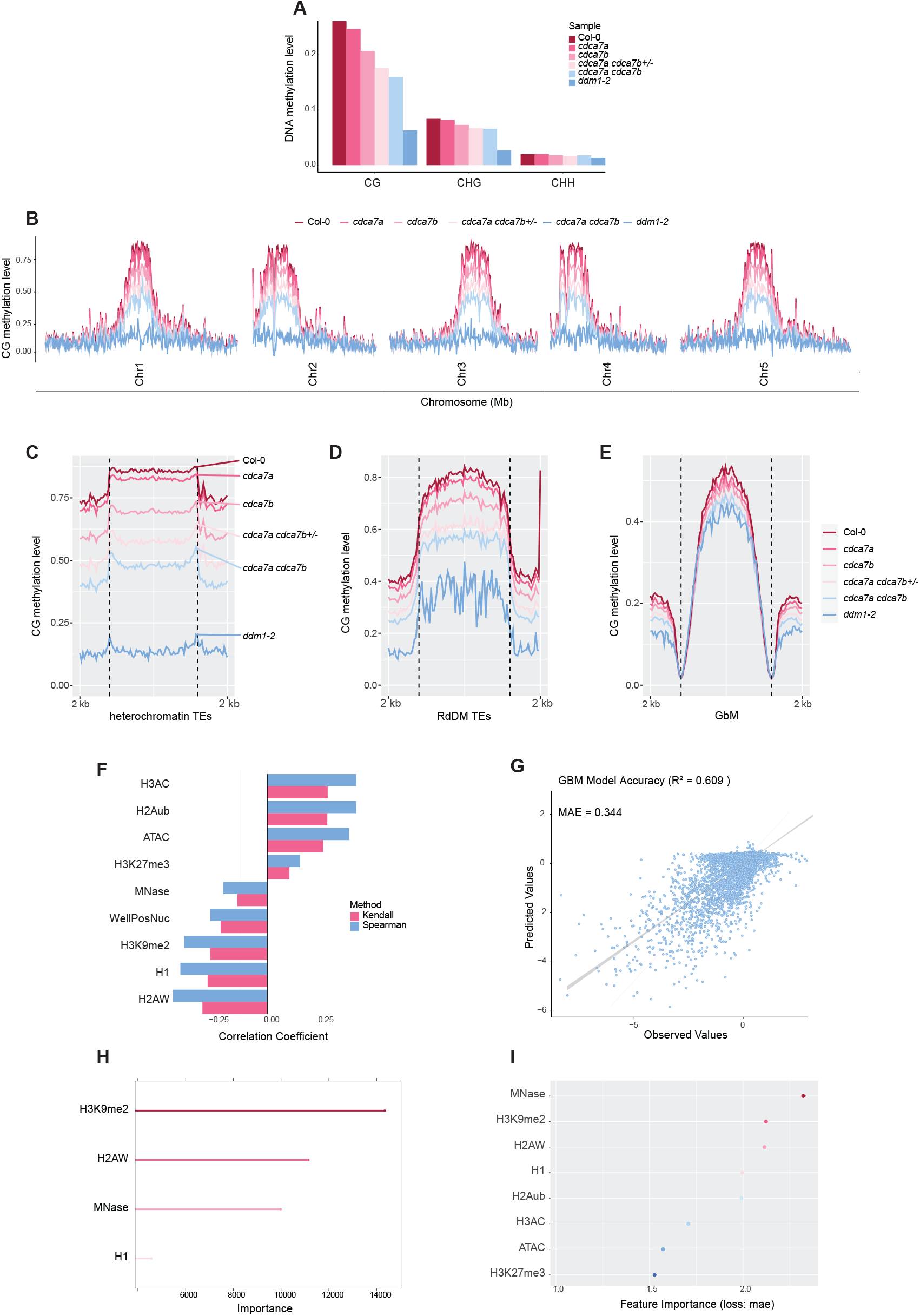
*CDCA7A* and *CDCA7B* maintain heterochromatic DNA methylation. **A**. Global DNA methylation summary of Col-0, *cdca7a, cdca7b, cdca7a cdca7b +/-, cdca7a cdca7b*, and *ddm1-2*. **B**. Genome-wide CG methylation landscape of Col-0, cdca7a, *cdca7b, cdca7a cdca7b+/-, cdca7a cdca7b*, and *ddm1-2*. Metaplots showing the CG methylation level at **C**. heterochromatin TEs, **D**. RdDM targeted TEs, and **E**. GbM across samples. **F**. Spearman and Kendall correlation coefficients between epigenetic features and CG methylation changes in the *cdca7a cdca7b* mutant compared to wild type. **G**. Prediction accuracy of generalized boosted regression model. **H**. Rank of the importance of epigenetic features via the generalized boosted regression model. **I**. The rank of the importance of epigenetic features, derived by shuffling each feature and measuring the reduction in performance.

Since *CDCA7A* and *CDCA7B* may be genetically redundant, we knocked out one *CDCA7B* allele in the *cdca7a* mutant background. Lacking one *CDCA7B* copy enhanced the *cdc7a* CG hypomethylation phenotype (Fig. 2A-E), while complete knockout of both *CDCA7A* and *CDCA7B* caused severe CG methylation loss at heterochromatic TEs (50% loss), compared to RdDM-targeted TEs (30% loss), and gene-body methylated (GbM) genes (10% loss). This reveals a genetic redundancy and a dosage-dependent role for *CDCA7B*. In contrast, CHG and CHH methylation decreased only slightly at heterochromatin (less than 10% loss) (Fig. S2B-C). Notably, *cdca7a cdca7b* mutants retained approximately 55% of wild-type pericentromeric CG methylation, which is much higher than the 10% in *ddm1-2* mutants and the 2% (near-complete loss) in *met1* mutants (Fig. 2B–E) [28]. Therefore, unlike CDCA7 in mammals, *CDCA7A* and *CDCA7B* are only partially required for DDM1 function *in vivo*. DDM1 may either remodel nucleosomes independently of *CDCA7A/B* or depend on additional methylation readers for recruitment and activity.

To further analyze the genome-wide patterns of DNA CG hypomethylation in *cdca7a cdca7b* mutants, we mapped hypo-methylated differentially methylated regions (hypoDMRs). These regions were mainly enriched in pericentromeric chromatin, which is characterized by high nucleosome density, H2A.W histone variants, H3K9me2 marks, and linker histone H1 (Fig. S2D-E). Supporting this, heterochromatic nucleosomes show about 27% CG methylation loss, while genic nucleosomes only exhibit a 3% reduction in *cdca7a cdca7b* (Fig. S2F-G). This suggests that heterochromatic nucleosomes preferentially depend on *CDCA7A* and *CDCA7B* for CG methylation maintenance.

To identify epigenetic features that predict CG methylation loss, we combined correlation analysis, machine learning, and feature importance ranking. Spearman and Kendall correlation coefficients showed strong links between *cdca7a cdc7b* CG hypomethylation and the heterochromatic features H2A.W, H1, and H3K9me2 (Fig. 2F, S2H). Conversely, transcription-activating marks like H3AC and H2Aub, along with ATAC-seq signals (indicating open chromatin) in wild-type, were associated with increased CG methylation in *cdca7a cdca7b* mutants (Fig. S2E, S2H). This increase in CG DNA methylation occurs at open chromatin regions, consistent with the redistribution of VIMs and MET1 to more accessible euchromatin when heterochromatin access is restricted due to the loss of *CDCA7A* and *CDCA7B*.

Using a generalized boosted regression model to predict CG methylation loss in *cdca7 cdca7b*, we achieved an *R*^*2*^ value of 0.609 and a mean absolute error of ∼0.3 (Fig. 2G). Among all features, H3K9me2, H2A.W, and nucleosome density (MNase-seq signal) emerged as the top predictors of CG hypomethylation in *cdca7 cdca7b* (Fig. 2H). To account for multicollinearity among features, we employed a permutation-based approach, shuffling individual features and quantifying their impact on model performance. Nucleosome density (MNase-seq) was the most critical predictor, followed by H3K9me2, H2A.W, and H1 enrichment (Fig. 2I). These findings demonstrate that *CDCA7A* and *CDCA7B* preferentially target heterochromatic nucleosomes marked by H3K9me2, H2A.W, and H1 for CG methylation maintenance. The interplay of these features underscores the chromatin context dependency of *CDCA7A/B* mediated CG methylation.

### *CDCA7A* and *CDCA7B* facilitate methyltransferase activity at H1-containing nucleosomes

DDM1 is essential for CG DNA methylation activity in H1-containing chromatin [6, 7]. To determine whether *CDCA7A/B* facilitates DNA methyltransferase activity at H1-containing nucleosomes, we generated *h1.1 h1.2 cdca7a cdca7b* quadruple mutants (Fig. S3A), and compared CG methylation levels to *those in cdca7a cdca7b* mutants. Genome-wide CG hypomethylation in *the cdca7a cdca7b* mutants was largely restored in the quadruple mutants, with DNA methylation levels nearly returning to wild-type at heterochromatic TEs, RdDM-targeted TEs, and GbM genes (Fig. 3A–F). These findings support that *CDCA7A/B* and DDM1 operate in the same pathway for DNA methylation maintenance, with one role being to counteract the repression of methylation caused by histone H1 enrichment.

**Figure 3.**
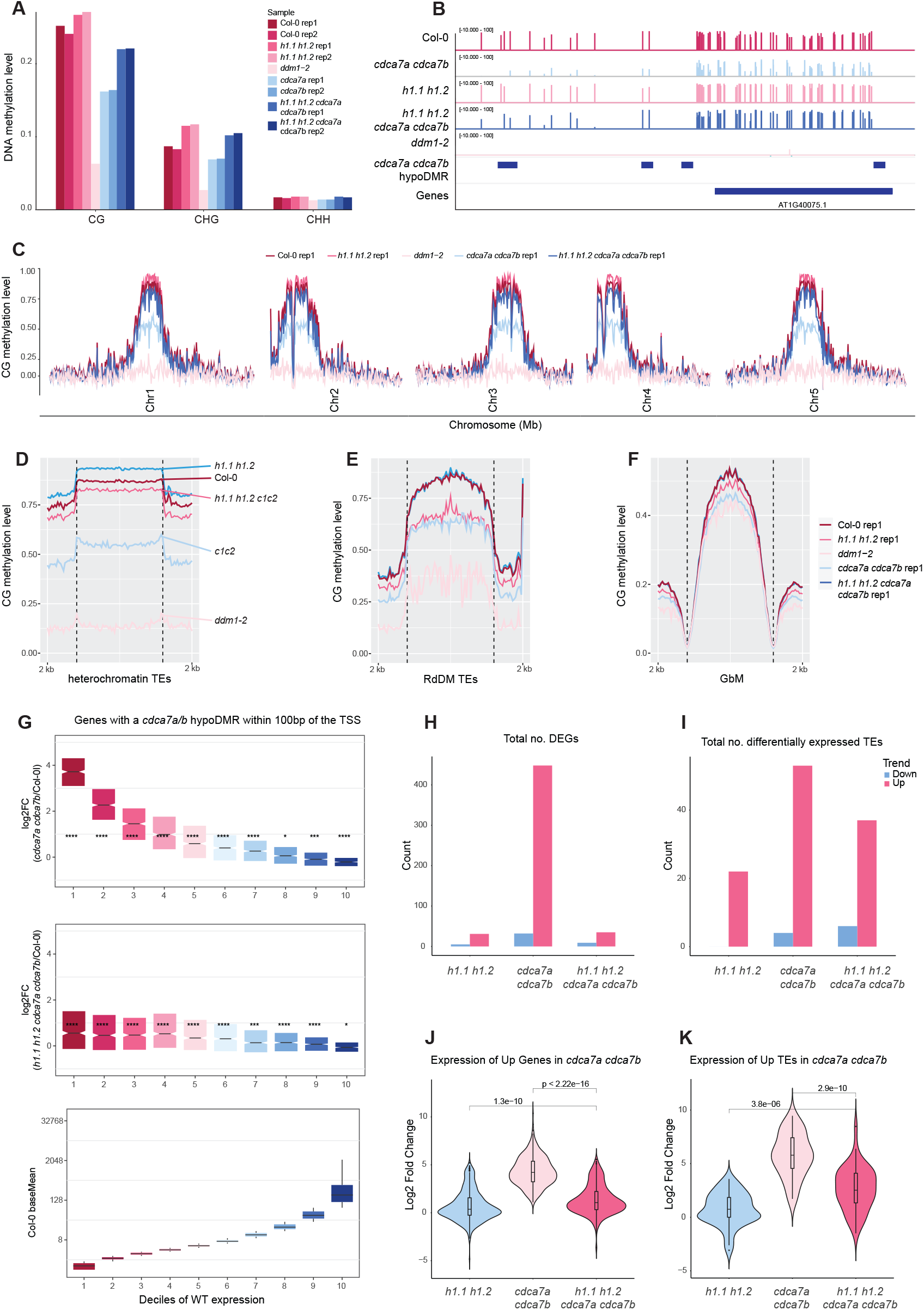
*CDCA7A* and *CDCA7B* facilitate methyltransferases to access H1-containing nucleosomes. **A**. Global DNA methylation summary of Col-0, *h1.1 h1.2, cdca7a cdca7b, h1.1 h1.2 cdca7a cdca7b*, and *ddm1-2*. **B**. Genome browser example showing CG methylation levels at the representative locus in Col-0, *cdca7a cdca7b, h1.1 h1.2, h1.1 h1.2 cdca7a cdca7b*, and *ddm1-2*. **C**. Genome-wide CG methylation landscape of Col-0, *h1.1 h1.2, cdca7a cdca7b, h1.1 h1.2 cdca7a cdca7b*, and *ddm1-2*. Metaplots showing the CG methylation level at **D**. heterochromatin TEs, **E**. RdDM targeted TEs, and **F**. GbM across samples. **G**. Normalized expression of genes proximal to the *cdca7a cdca7b* hypoDMR sites. The bottom bar plot shows the ten deciles clustered by wild-type expression level. Total number of differentially expressed **H**. genes, and **I**. TEs in *h1.1 h1.2, cdca7a cdca7b*, and *h1.1 h1.2 cdca7a cdca7b*. Log2 fold changes of expression of upregulated **J**. genes, and **K**. TEs in the *cdca7a cdca7b* mutant.

We found that a subset of hypoDMRs, located at heterochromatin-euchromatin boundaries, remained DNA hypomethylated in *h1.1 h1.2 cdca7a cdca7b* mutants (Fig. S3B). These regions showed lower nucleosome density, reduced chromatin accessibility, and depletion of repressive histone marks (Fig. S3C-F), indicating that *CDCA7A* and *CDCA7B* target inaccessible heterochromatin edges. We also found that MORC proteins, ATP-dependent enzymes that compact and silence chromatin (Moissiard, 2012 #58), preferentially target regions that remain hypomethylated in *h1.1 h1.2 cdca7a cdca7b* mutants (Fig. S3G). Therefore, outside of H1-enriched nucleosomes, *CDCA7A* and *CDCA7B* may assist VIM-MET1 activity to overcome MORC-mediated chromatin compaction. CG hypomethylation at *cdca7a cdca7b* hypoDMRs triggered mRNA upregulation of proximal genes (within 100 bp), especially those with low baseline expression in wild type (Fig. 3G). Notably, H1 depletion restored the expression of most *cdca7a cdca7b* upregulated genes to near-wild type levels (Fig. 3G). Beyond the genes proximal to *cdca7a cdca7b* hypoDMRs, genome-wide, *cdca7a cdca7b* mutants activated approximately 450 genes, with most (85%) becoming transcriptionally silent in *h1.1 h1.2 cdca7a cdca7b* (Fig. 3H). Additionally, about 55 TEs became significantly activated in *cdca7a cdca7b* (Fig. 3I). A subset of the *cdca7a cdca7b* activated TEs became downregulated by H1 loss, while TEs close to the remaining *h1.1 h1.2 cdca7a cdca7b* hypoDMRs remained upregulated compared to wild type (Fig. 3I, Fig. S3H).

When examining the extent of gene and TE transcript upregulation, the loss of H1 significantly reduced the degree of upregulation at activated genes, and also partially at activated TEs (Fig. 3J-K), consistent with the degree of DNA methylation rescue. These findings demonstrate that *CDCA7A* and *CDCA7B* mainly operate in H1-enriched heterochromatin to preserve DNA methylation and enforce transcriptional silencing. Their ability to act at heterochromatin boundaries, independent of H1, also highlights a context-specific role.

### *CDCA7A* and *CDCA7B* promote DNA methylation at centromeres

Since ICF syndrome-associated *cdca7* mutations cause DNA hypomethylation at human centromere alpha-satellite repeats [15], we hypothesized that *Arabidopsis CDCA7A* and *CDCA7B* might similarly regulate repetitive regions, including centromeric *CEN178* satellite repeats. To test this, we analyzed DNA methylation at centromeric regions using the Col-CEN-v1.2 genome assembly [16], which fully resolves centromeric sequences. Loss of *CDCA7A* and *CDCA7B* resulted in approximately a 50% reduction in CG methylation at satellite repeats, while non-CG methylation was only modestly affected (15%) (Fig. 4A, Fig. S4A-B). Within the *CEN178* repeat arrays, not only was CG methylation substantially reduced, but its distribution also shifted toward linker DNA regions (Fig. 4B-D). This pattern suggests that *CDCA7A* and *CDCA7B* are required for proper methylation distribution at CENH3-containing nucleosomes. Supporting this, the decrease in CG methylation was more pronounced at *CEN178* repeats with higher CENH3 enrichment (Fig. 4E-F).

**Figure 4.**
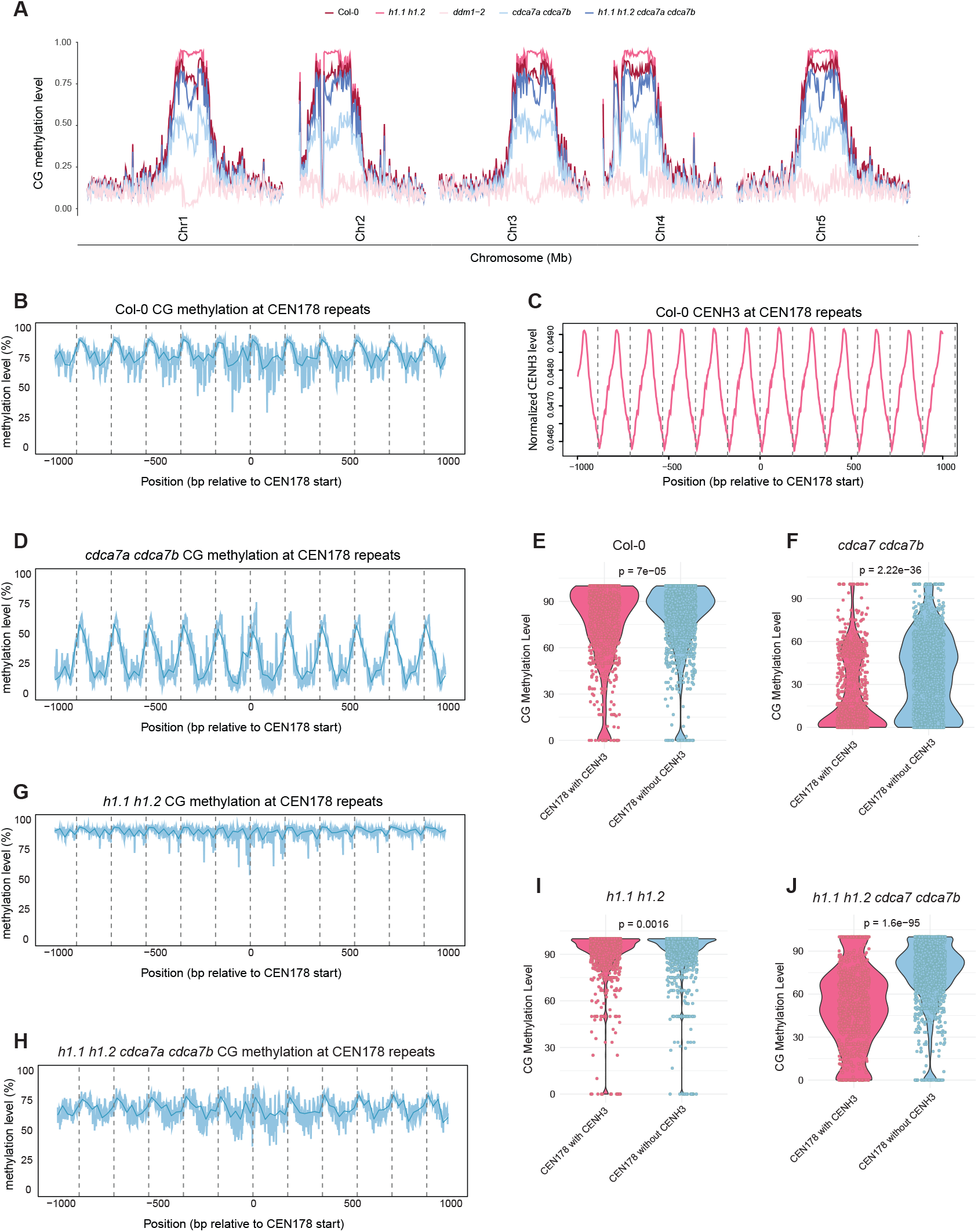
*CDCA7A* and *CDCA7B* promote CG methylation establishment at centromeres independently of H1. **A**. Global CG methylation summary of Col-0, *h1.1 h1.2, cdca7a cdca7b, h1.1 h1.2 cdca7a cdca7b*, and *ddm1-2*, using Col-Cen-v1.2 as the reference genome. **B**. Metaplot showing the CG methylation level at CEN178 satellite repeats in Col-0. **C. Metaplot showing the normalized wild type CENH3 enrichment at the CEN178 satellite repeats. D**. Metaplot showing the CG methylation level at CEN178 satellite repeats in the *cdca7a cdca7b* mutant. Violin plots showing the CG methylation level at CEN178 satellite repeats with and without CENH3 enrichment in **E**. Col-0, and **F**. *cdca7a cdca7b*. Metaplots showing the CG methylation level at CEN178 satellite repeats in **G**. *h1.1 h1.2* and **H**. *h1.1 h1.2 cdca7a cdca7*. Violin plots showing the CG methylation level at CEN178 satellite repeats with and without CENH3 enrichment in **I**. *h1.1 h1.2*, and **J**. *h1.1 h1.2 cdca7a cdca7b* mutants.

We next assessed whether H1 depletion in the *cdca7a cdca7b* mutant background could similarly restore DNA methylation loss, as we previously observed in the chromosome arms and pericentromeres. Compared to *cdca7a cdca7b* mutants, *h1.1 h1.2 cdca7a cdca7b* quadruple mutants gained CG methylation within the *CEN178* satellite arrays, but only partially (60% versus 80% in wild-type), indicating that *CDCA7A* and *CDCA7B* function beyond H1 in the centromeres (Fig. 4A). This aligns with the incomplete rescue of DNA methylation at heterochromatin boundaries seen in *h1.1 h1.2 cdca7a cdca7b* mutants (Fig. S3B).

To directly examine the role of *CDCA7A* and *CDCA7B* in centromeric CG methylation without H1, we compared *h1.1 h1.2* and *h1.1 h1.2 cdca7a cdca7b* mutants. Strikingly and unexpectedly, CG methylation, and to a lesser extent, non-CG methylation, increased significantly with H1 loss (Fig. 4A, S4A-B). At *CEN178* satellite arrays, CG methylation approaches nearly 100% in *h1.1 h1.2*, indicating that H1 acts as a barrier to DNA methylation in *Arabidopsis* centromeres (Fig. 4A-C, 4G). In the *h1.1 h1.2* mutants, CG methylation was primarily gained in the centers of CEN178 repeats, which are typically enriched for CENH3 (Fig. 4B-C, 4G). This CG hypermethylation in the middle of the *CEN178* repeats was lost in the *h1.1 h1.2 cdca7a cdca7b* mutants (Fig. 4C, 4G-H). Consistently, the CG methylation decrease in the quadruple mutant was more pronounced within CENH3-enriched *CEN178* repeats, further emphasizing the role of *CDCA7A* and *CDCA7B* in maintaining DNA methylation at centromeric nucleosomes (Fig. 4I-J). However, the increase in CHG and CHH DNA methylation in *h1.1 h1.2* was unaffected by the loss of *CDCA7A and CDCA7B* (Fig. S4A-F), indicating that they specifically promote VIM and MET-mediated CG methylation.

We propose that the depletion of histone H1 broadly decompacts heterochromatin, allowing increased access for *CDCA7A* and *CDCA7B* to the centromeres thereby promoting CG hypermethylation by VIM and MET1 through a replication-uncoupled mechanism, to compensate for imperfect replication-coupled maintenance [15, 18]. However, the mechanism by which *CDCA7A* and *CDCA7B* are recruited to centromeres remains unclear. One possibility is that the conserved zf-CXXC_R1 domain in *CDCA7A* and *CDCA7B* recognizes non-B DNA structures in centromeric satellites, as shown *in vitro* [17]. Alternatively, *CDCA7A* and *CDCA7B* might directly recognize satellite repeats *CEN178*, or associated centromeric chromatin marks, diverging from the binding preferences seen in mammalian CDCA7.

## Discussion

This work indicates that *CDCA7A* and *CDCA7B*, the *Arabidopsis* counterparts of mammalian CDCA7, interact with DDM1 to help maintain CG methylation. We propose that *CDCA7A* and *CDCA7B* promote VIM and MET1 access to tightly packed heterochromatin by remodeling H1-containing nucleosomes at pericentromeric regions. This idea is supported by the rescue of pericentromeric DNA hypomethylation in *cdca7a cdca7b* mutants when H1 is lost in *h1.1 h1.2 cdca7a cdca7b* quadruple mutants. We also discovered a role for *CDCA7A* and *CDCA7B* in promoting centromeric CG methylation in the absence of H1. We suggest that CDCA7A and *CDCA7B* remodel CENH3-containing nucleosomes and provide better access to methyltransferases, including but not limited to VIM and MET1.

Importantly, our study reveals a novel role for H1 in preventing DNA methylation at centromeric chromatin. Since the loss of H1 results in nearly saturated CG methylation at satellite repeats, primarily due to *CDCA7A* and *CDCA7B* activity, we suggest that H1-bound hemi-methylated nucleosomes are common features of *Arabidopsis* centromeres. H1 depletion then allows DNA methylation maintenance through the CDCA7 pathway. This discovery provides insight into the mechanism behind the low DNA methylation levels at mammalian centromeres, with a high abundance of linker histone H1 likely restricting centromeric DNA methylation. *CDCA7A* and *CDCA7B* may directly recognize centromeric DNA motifs to facilitate methylation establishment and maintenance in these regions. The conserved zf-CXXC_R1 domain of *CDCA7A* and *CDCA7B* can bind both canonical and non-B DNA structures [17], but its exact targeting preferences remain unknown. Future studies combining *in vitro* binding assays and *in vivo* mutagenesis will clarify whether centromeric repeats or structural features determine *CDCA7A/CDCA7B* localization.

While *cdca7a cdca7b* mutants show DNA hypomethylation, the extent of reduction is less than in *ddm1-2* and *met1* mutants. This difference in phenotypic severity suggests two possibilities: (1) *Arabidopsis* Class II/III CDCA7 homologs might partially compensate for *CDCA7A* and *CDCA7B* in recruiting DDM1, or (2) DDM1 can still partially localize to heterochromatin without CDCA7 proteins. Further research into DDM1 recruitment mechanisms is needed to clarify this. These findings underscore the conserved function of CDCA7 proteins in linking chromatin remodelers and methyltransferases across different species, while also revealing plant-specific changes in centromere regulation. By clarifying how *CDCA7A/CDCA7B* control H1-dependent and H1-independent methylation, this work enhances our understanding of heterochromatin and DNA methylation dynamics.

## Methods and Materials

### Phylogenetic Analysis

Highly conserved zf-CXXC_R1 domain sequences of *Arabidopsis* Class I (CDCA7A and CDCA7B), Class II (AT1G67270, AT1G67780, and AT5G38690), Class III (AT1G09060,

AT1G11950, and AT3G07610) CDCA7 and human CDCA7 were taken for phylogenetic analysis. All the sequences were listed in Figure 1C. Protein sequence alignments were performed using Clustal Omega. A graphic representation of the phylogenetic tree was generated using Jalview.

### AlphaFold prediction

Full-length proteins of *CDCA7A/B* and DDM1 are taken for AlphaFold3 prediction [27]. For each prediction, the best model was selected for further structural analysis. A cutoff distance of 5 Å was applied. The protein structures were visualized using Pymol.

### Plant materials and growth conditions

All plants used in this paper were *Arabidopsis thaliana* Col-0 ecotype and were grown under long-day conditions (16 h light and 8 h dark). Seedlings of the Col-0, *cdca7a, cdca7b, cdca7a cdca7b+/-, cdca7a cdca7b, h1.1 h1.2, h1.1 h1.2 cdca7a cdca7b*, and *ddm1-2* were harvested after 14-day incubation under long-day conditions. The T-DNA insertion lines used in this study are listed here: h1.1 (SALK_128430C), and h1.2 (GABI_406H11_012502). *ddm1-2* contains a splice donor site mutation. CRISPR mutants were generated using the pBEE401E CRISPR system [29]. CDCA7A CRISPR mutant was generated using two combinations of guides: GGGGTTTCTTTGATTAGTTC and TTGGGAATACAGAAAGAAGC or CAAAGGTCTCTCTTTACGAA and ATCCCATCAGTGTAGATAAC. CDCA7B CRISPR mutant was generated using two combinations of guides: TTCGCTCTCGTTCTCACCAC and AAGGCCAGAGATTTACACTG or GAGTTTCCTCCTCCGACTGT and GGTTCCTCTGCGTAGGAAAC.

### WGBS

14-day old seedlings were harvested from Col-0, *cdca7a, cdca7b, cdca7a cdca7b+/-, cdca7a cdca7b, h1.1 h1.2, h1.1 h1.2 cdca7a cdca7b*, and *ddm1-2*. Samples were immediately frozen in liquid nitrogen. DNA was extracted using the DNeasy Plant Mini kit (Qiagen). A total of 100ng DNA was sheared to ∼200bp using the Covaris S2 (Covaris). Then the libraries were constructed using the Ovation Ultralow Methyl-seq kit (NuGEN), and bisulfite conversion was achieved using the Epitect Bisulfite Conversion kit (QIAGEN). Finally, the libraries were sequenced on Illumina Novaseq X plus instruments.

### Correlation analysis

Epigenetic data were downloaded from published datasets. H1, H3K9me2, MNase-seq, H3AC, H2Aub, H3K27me3, H2A.W, and ATAC-seq were normalized using the preprocess function from the caret library. Then, correlation coefficients were calculated between variables using the Spearman and Kendall algorithms, provided by the corrplot package. Visualizations of the correlation matrix were performed using the ggplot2 package.

### Machine learning

Epigenetic features with high correlation (>0.78) were removed before data normalization. 75% of data points were used for training, and the remaining 25% were used for testing. The Gradient Boosted Regression (GBM) model was selected based on its highest performance. A 10-fold cross-validation was repeated 10 times to examine model performance. Tuning parameters were applied as follows: interaction depth (3,6,9), number of trees (200, 250, 300). Finally, R-squared and MAE values were calculated, and the importance of epigenetic features was ranked.

### Feature importance ranking

Feature importance measure was alternatively performed using the iml package. Each feature was shuffled and the decrease in the GBM model performance was calculated. The loss in performance was measured with MAE.

### RNA-seq

Three biological replicates were used for each genotype. An individual 2-week-old seedling was collected as a biological replicate and frozen in liquid nitrogen. The samples were then ground into powder, and RNA was extracted using the Direct-zol RNA MiniPrep kit (Zymo Research). 500ng of total RNA from seedling tissues were used for RNA-seq library preparation with TruSeq Stranded mRNA kit (Illumina). The final library was sequenced on Illumina Novaseq X plus instruments.

### WGBS analysis

WGBS were filtered and Illumina adaptors were removed using Trim Galore (v 0.6.7, Babraham Institute). Reads with three or more consecutively methylated CHH sites were treated as non-converted reads and filtered out. Bismark (v 0.19.1, Babraham Institute) [30] was used to map the reads to the Arabidopsis reference genome (TAIR10) and Col-Cen-v1.2 genome assembly [23]. ViewBS (v 0.1.11) was used to generate the plots [31]. DNA methylation levels across 1 kb upstream and downstream of CEN178 centers were first computed using deepTools (v 3.0.2) (Ramirez, 2016 #54) with the option of computeMatrix reference-point. The output matrix was then processed with in-house Perl and R scripts to generate the final methylation profile plots.

### RNA-seq analysis

RNA-seq reads were filtered and Illumina adaptors were trimmed using Trim Galore (v 0.6.7, Babraham Institute). Left reads were mapped to the Arabidopsis reference genome (TAIR10) using STAR (v 2.7.11a) [32]. Uniquely mapped reads with less than 5% of mismatches were kept. For visualization, bigwig files were generated using deeptools (v 3.0.2) [33] bamCoverage with the options -- normalizeUsing RPGC and --binSize 10. HTSeq (v 0.13.5) was used to obtain the read counts for genes and TE. DESeq2 (v 1.42.0). Differential analysis was used perform via DESeq2 (v 1.42.0) with the cutoff padj < 0.05 and |log2FC| >= 1. Eventually, ggplot2 (v 3.4.4) was sued to generate all the related plots.

## Data availability

The high-throughput sequencing data generated in this paper will be deposited in the Gene Expression Omnibus (GEO) database.

## Acknowledgments

We thank Dr. Colette Picard and Dr. Zhongshou Wu for their discussion and advice. We also thank Mahnaz Akhavan and the UCLA BSCRC BioSequencing Core for the sequencing support. This work was supported by S.E.J. funding from the Howard Hughes Medical Institute.

## Authors Contributions Statement

S.W., S.E.J., and I.R.H. conceived the study, designed the research, and wrote the manuscript. S.W. performed most of the experiments and data analysis. T.L. and M.N. contributed to the data analysis. R.C., E.K.L., C.F., and Y.H. contributed to the experiments. S.F. performed BS-PCR-seq and all high-throughput sequencing.

## Figure Legends

**Supplementary Figure 1.**
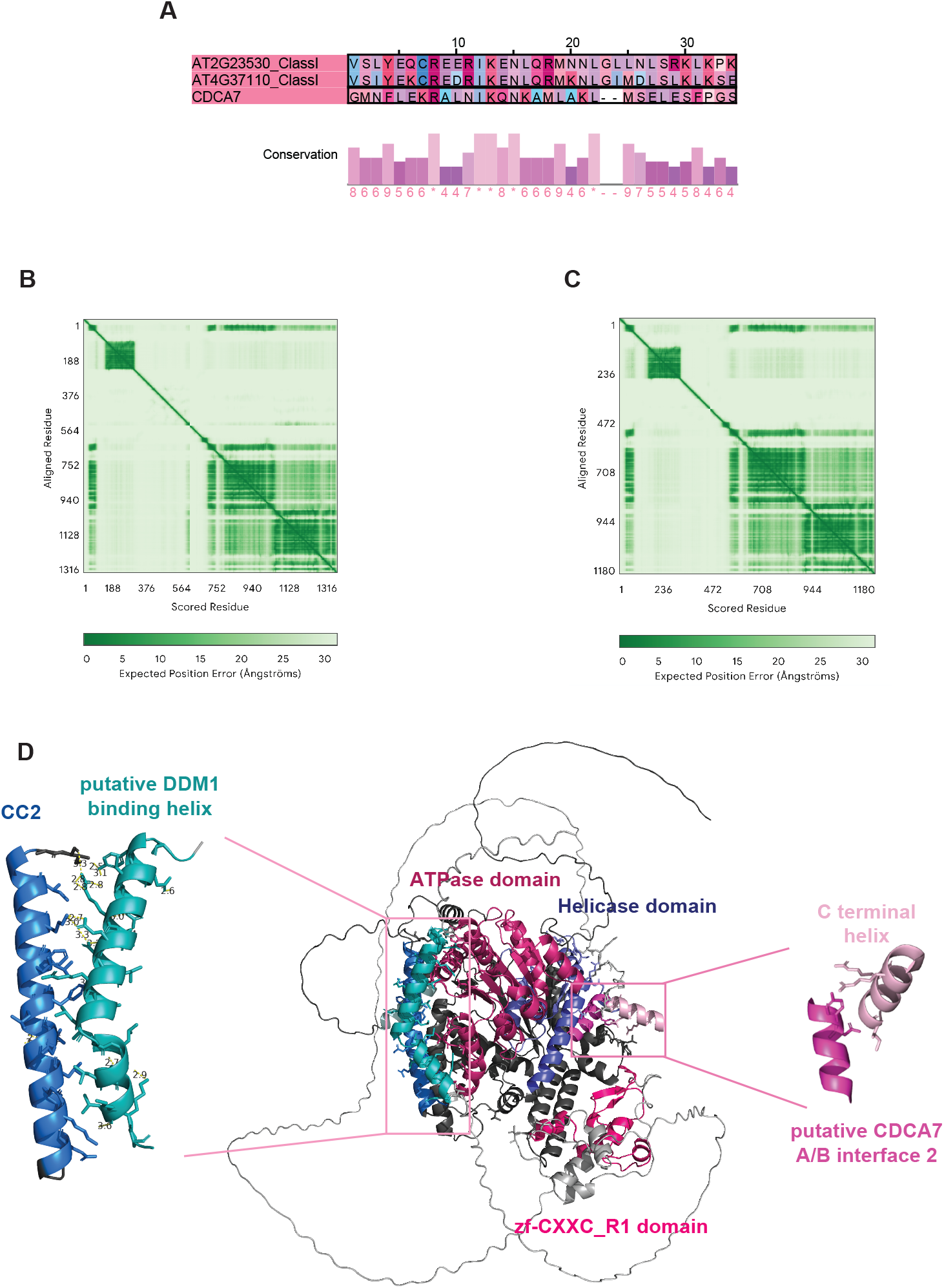
*CDCA7A* and *CDCA7B* retain the HLBH domain for interacting with DDM1. **A**. Sequence alignment calculated by Clustal Omega of the HLBH domains of *homo sapiens* CDCA7 and *Arabidopsis* Class I homologs. The red asterisk indicated the point mutations. Predicted Aligned Error (PAE) from AF3 structural modeling of **B**. *CDCA7A* or **C**. *CDCA7B* and DDM1 interactions. **D**. Close illustration of the AF3 predicted interface between *CDCA7B* and DDM1. Cyan represents the putative DDM1-binding helix of *CDCA7B*. Blue represents CC2 of DDM1. Light pink represents the C-terminal helix of *CDCA7B*. Pink represents putative CDCA7 interface 2 of DDM1. Dark pink represents the ATPase domain of DDM1. Purple indicates the Helicase C term domain of DDM1. Bright pink represents the zf-CXXC_R1 domain of *CDCA7B*.

**Supplementary Figure 2.**
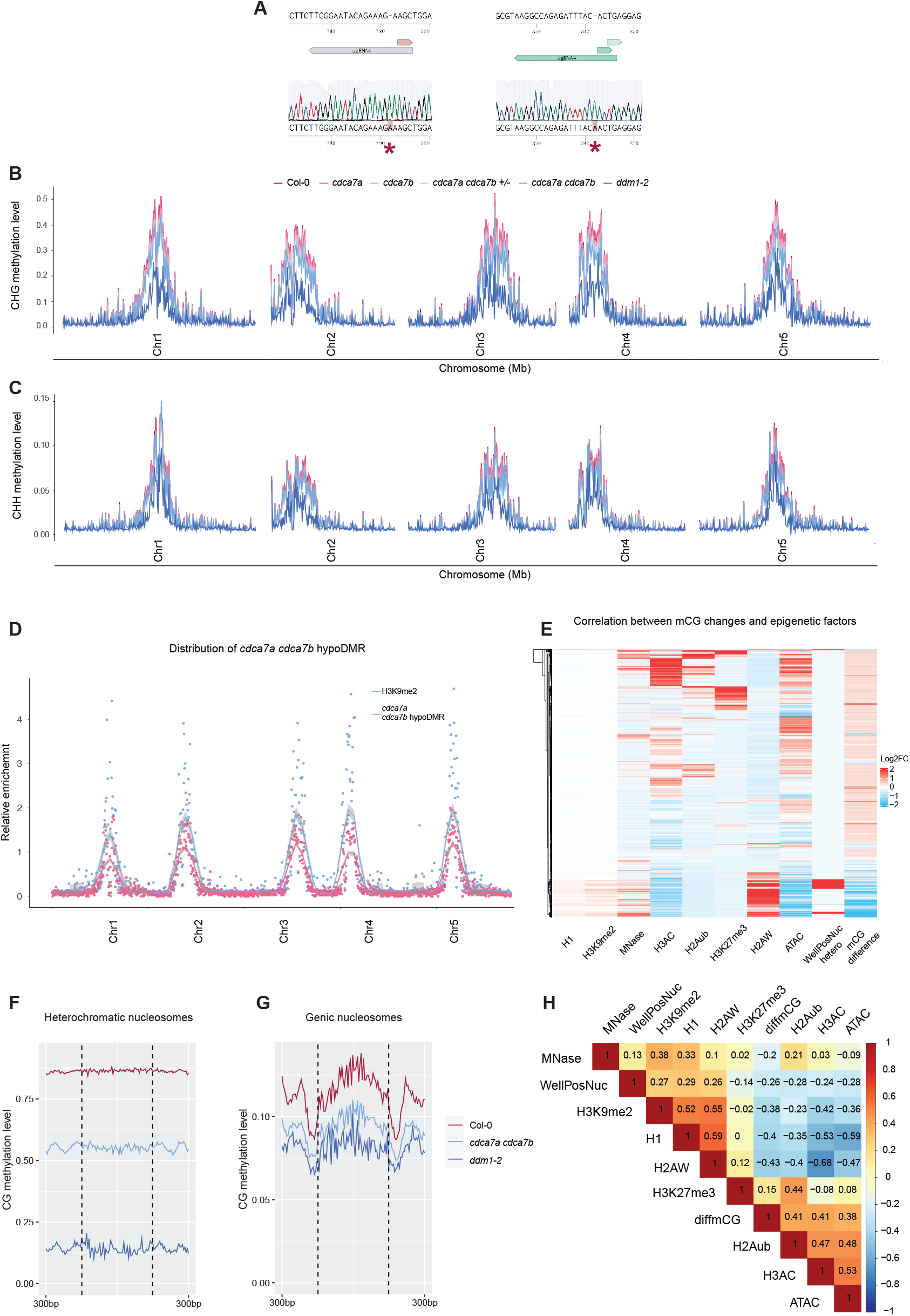
*CDCA7A* and *CDCA7B* promote DNA methylation at heterochromatic nucleosomes. **A**. Sanger sequencing confirmation of CRISPR-Cas9 introduced A insertion at the *CDCA7A* and *CDCA7B* coding regions. **B**. Genome-wide CHG methylation landscape of Col-0, *cdca7a, cdca7b, cdca7a cdca7b+/-, cdca7a cdca7b*, and *ddm1-2*. **C**. Genome-wide CHH methylation landscape of Col-0, *cdca7a, cdca7b, cdca7a cdca7b+/-, cdca7a cdca7b*, and *ddm1-2*. **D**. Distribution of the *cdca7a cdca7b* hypoDMR. H3K9me2 enrichment marks the locations of heterochromatin. **E**. Heatmaps showing the relation between epigenetic features and CG methylation changes in the *cdca7a cdca7b* mutant. Metaplots showing CG methylation levels at **F**. heterochromatic well-positioned nucleosomes and **G**. genic well-positioned nucleosomes of Col-0, *cdca7a cdca7b*, and *ddm1-2*. **H**. Spearman correlation matrix among epigenetic features and changes in CG methylation level in the *cdca7a cdca7b* mutants.

**Supplementary Figure 3.**
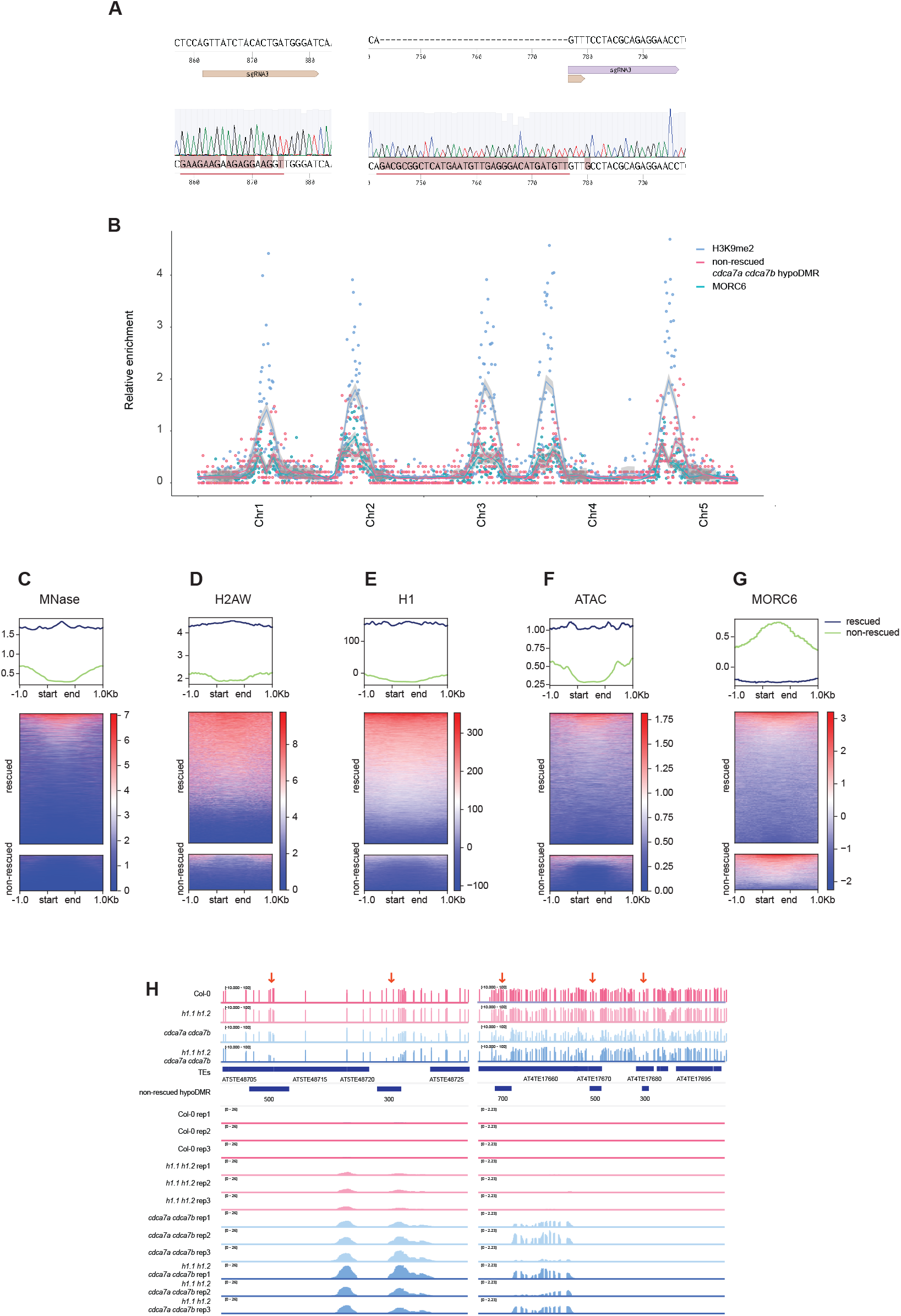
*CDCA7A* and *CDCA7B* maintain CG methylation independently of H1. **A**. Sanger sequencing confirmation of CRISPR-Cas9 introduced mutations at *CDCA7A* and *CDCA7B* coding regions. The red lines indicated the region where mutations were introduced. **B**. Distribution of the non-rescued *cdca7a cdca7b* hypoDMR (regions with CG methylation levels not recovered in *h1.1 h1.2 cdca7a cdca7b*). Relative enrichment of H3K9me2 indicates heterochromatin. Metaplots showing **C**. MNase-seq signal, **D**. H2A.W ChIP-seq signal, **E**. H1 ChIP-seq signal, **F**. ATAC-seq signal, and **G**. MORC6 ChIP-seq signal at non-rescued *cdca7a cdca7b* hypoDMR sites. **H**. Genome browser examples showing CG methylation and gene expression across Col-0, *h1.1 h1.2, cdca7a cdca7b*, and *h1.1 h1.2 cdca7a cdca7b* at non-rescued hypoDMRs in *h1.1 h1.2 cdca7a cdca7b* mutants. The red arrows indicate the locations of the hypoDMRs in the *h1.1 h1.2 cdca7a cdca7b* mutants.

**Supplementary Figure 4.**
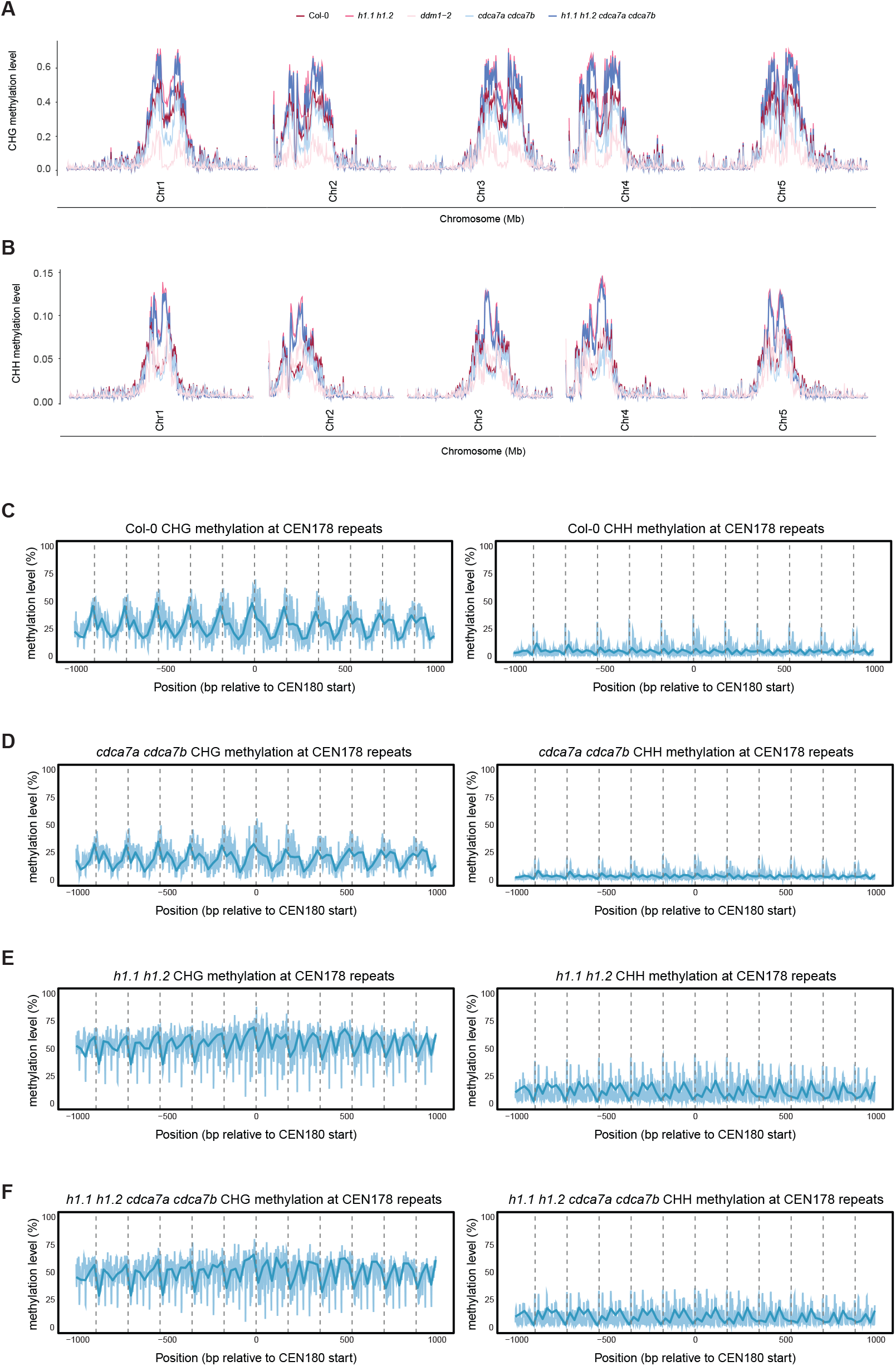
*CDCA7A* and *CDCA7B* play a minor role in regulating non-CG methylation at centromeric regions. Genome-wide **A**. CHG methylation, and **B**. CHH methylation landscapes of Col-0, *h1.1 h1.2, cdca7a cdca7b, h1.1 h1.2 cdca7a cdca7b*, and *ddm1-2*. Metaplots showing the non-CG methylation levels at CEN178 satellite repeats of **C**. Col-0, **D**. *cdca7a cdca7b*, **E**. *h1.1 h1.2*, and **F**. *h1.1 h1.2 cdca7a cdca7b*.

**Supplementary Table 1. Details of the predicted interaction interfaces between *CDCA7A* and DDM1 from AF3**.

**Supplementary Table 2. Details of the predicted interaction interfaces between *CDCA7B* and DDM1 from AF3**.

## Reference

1. Zhang, H., Z. Lang, and J.K. Zhu, Dynamics and function of DNA methylation in plants. Nat Rev Mol Cell Biol, 2018. 19(8): p. 489–506.

2. Smith, Z.D. and A. Meissner, DNA methylation: roles in mammalian development. Nat Rev Genet, 2013. 14(3): p. 204–20.

3. Law, J.A. and S.E. Jacobsen, Establishing, maintaining and modifying DNA methylation patterns in plants and animals. Nat Rev Genet, 2010. 11(3): p. 204–20.

4. Stroud, H., et al., Non-CG methylation patterns shape the epigenetic landscape in Arabidopsis. Nat Struct Mol Biol, 2014. 21(1): p. 64–72.

5. Cao, X., et al., Role of the DRM and CMT3 methyltransferases in RNA-directed DNA methylation. Curr Biol, 2003. 13(24): p. 2212–7.

6. Zemach, A., et al., The Arabidopsis nucleosome remodeler DDM1 allows DNA methyltransferases to access H1-containing heterochromatin. Cell, 2013. 153(1): p. 193–205.

7. Lyons, D.B. and D. Zilberman, DDM1 and Lsh remodelers allow methylation of DNA wrapped in nucleosomes. Elife, 2017. 6.

8. Choi, J., et al., DNA Methylation and Histone H1 Jointly Repress Transposable Elements and Aberrant Intragenic Transcripts. Mol Cell, 2020. 77(2): p. 310–323 e7.

9. Harris, C.J., et al., H1 restricts euchromatin-associated methylation pathways from heterochromatic encroachment. Elife, 2024. 12.

10. Osakabe, A., et al., Molecular and structural basis of the chromatin remodeling activity by Arabidopsis DDM1. Nat Commun, 2024. 15(1): p. 5187.

11. Lee, S.C., et al., Chromatin remodeling of histone H3 variants by DDM1 underlies epigenetic inheritance of DNA methylation. Cell, 2023. 186(19): p. 4100–4116 e15.

12. Osakabe, A., et al., The chromatin remodeler DDM1 prevents transposon mobility through deposition of histone variant H2A.W. Nat Cell Biol, 2021. 23(4): p. 391–400.

13. Jenness, C., et al., HELLS and CDCA7 comprise a bipartite nucleosome remodeling complex defective in ICF syndrome. Proc Natl Acad Sci U S A, 2018. 115(5): p. E876– E885.

14. Funabiki, H., et al., Coevolution of the CDCA7-HELLS ICF-related nucleosome remodeling complex and DNA methyltransferases. Elife, 2023. 12.

15. Wassing, I.E., et al., CDCA7 is an evolutionarily conserved hemimethylated DNA sensor in eukaryotes. Sci Adv, 2024. 10(34): p. eadp5753.

16. Shinkai, A., et al., The C-terminal 4CXXC-type zinc finger domain of CDCA7 recognizes hemimethylated DNA and modulates activities of chromatin remodeling enzyme HELLS. Nucleic Acids Res, 2024. 52(17): p. 10194–10219.

17. Hardikar, S., et al., The ICF syndrome protein CDCA7 harbors a unique DNA binding domain that recognizes a CpG dyad in the context of a non-B DNA. Sci Adv, 2024. 10(34): p. eadr0036.

18. Han, M., et al., A role for LSH in facilitating DNA methylation by DNMT1 through enhancing UHRF1 chromatin association. Nucleic Acids Res, 2020. 48(21): p. 12116–12134.

19. Tachiwana, H., et al., Crystal structure of the human centromeric nucleosome containing CENP-A. Nature, 2011. 476(7359): p. 232–5.

20. Thakur, J., J. Packiaraj, and S. Henikoff, Sequence, Chromatin and Evolution of Satellite DNA. Int J Mol Sci, 2021. 22(9).

21. Logsdon, G.A., et al., The variation and evolution of complete human centromeres. Nature, 2024. 629(8010): p. 136–145.

22. Altemose, N., et al., Complete genomic and epigenetic maps of human centromeres. Science, 2022. 376(6588): p. eabl4178.

23. Naish, M., et al., The genetic and epigenetic landscape of the Arabidopsis centromeres. Science, 2021. 374(6569): p. eabi7489.

24. Wlodzimierz, P., et al., Cycles of satellite and transposon evolution in Arabidopsis centromeres. Nature, 2023. 618(7965): p. 557–565.

25. Hou, X., et al., A near-complete assembly of an Arabidopsis thaliana genome. Mol Plant, 2022. 15(8): p. 1247–1250.

26. Brzeski, J. and A. Jerzmanowski, Deficient in DNA methylation 1 (DDM1) defines a novel family of chromatin-remodeling factors. J Biol Chem, 2003. 278(2): p. 823–8.

27. Abramson, J., et al., Accurate structure prediction of biomolecular interactions with AlphaFold 3. Nature, 2024. 630(8016): p. 493–500.

28. Kankel, M.W., et al., Arabidopsis MET1 cytosine methyltransferase mutants. Genetics, 2003. 163(3): p. 1109–22.

29. Wang, Z.P., et al., Egg cell-specific promoter-controlled CRISPR/CasS efficiently generates homozygous mutants for multiple target genes in Arabidopsis in a single generation. Genome Biol, 2015. 16(1): p. 144.

30. Krueger, F. and S.R. Andrews, Bismark: a flexible aligner and methylation caller for Bisulfite-Seq applications. Bioinformatics, 2011. 27(11): p. 1571–2.

31. Huang, X., et al., ViewBS: a powerful toolkit for visualization of high-throughput bisulfite sequencing data. Bioinformatics, 2018. 34(4): p. 708–709.

32. Dobin, A., et al., STAR: ultrafast universal RNA-seq aligner. Bioinformatics, 2013. 29(1): p. 15–21.

33. Ramirez, F., et al., deepTools2: a next generation web server for deep-sequencing data analysis. Nucleic Acids Res, 2016. 44(W1): p. W160–5.

