## Supplementary Tables for "CDCA7 facilitates MET1-mediated CG DNA methylation maintenance in centromeric heterochromatin via histone H1"

**Supplementary Table 1: CDCA7A DDM1 interaction Residues**

| residue_1 | residue_2 | distance | chain_1 | chain_2 | interaction |
|---|---|---|---|---|---|
| A:ARG:35: | B:ASN:356: | 2.42 | A | B | Polar |
| A:ARG:35: | B:ASP:355: | 4.02 | A | B | Contact |
| A:ARG:35: | B:GLN:218: | 3.52 | A | B | Contact |
| A:ARG:35: | B:GLY:220: | 4.2 | A | B | Contact |
| A:ARG:35: | B:TRP:217: | 2.97 | A | B | HBond,Polar |
| A:ARG:38: | B:ASN:219: | 4.32 | A | B | Contact |
| A:ARG:38: | B:GLN:218: | 2.84 | A | B | Polar |
| A:ARG:38: | B:GLU:132: | 2.76 | A | B | Polar |
| A:ARG:38: | B:GLY:128: | 4.95 | A | B | Contact |
| A:ARG:38: | B:ILE:124: | 3.84 | A | B | Contact |
| A:ARG:38: | B:ILE:129: | 3.34 | A | B | Polar |
| A:ARG:45: | B:PHE:116: | 3.61 | A | B | Contact |
| A:ARG:45: | B:TYR:113: | 4.42 | A | B | Contact |
| A:ARG:547: | B:LEU:512: | 3.27 | A | B | Contact |
| A:ARG:547: | B:TYR:511: | 3.54 | A | B | Contact |
| A:ARG:547: | B:TYR:513: | 3.97 | A | B | Contact |
| A:ARG:547: | B:TYR:558: | 4.31 | A | B | Contact |
| A:ARG:549: | B:ARG:569: | 4.96 | A | B | Contact |
| A:ARG:549: | B:ASP:554: | 4.9 | A | B | Contact |
| A:ARG:549: | B:ASP:571: | 4.13 | A | B | Contact |
| A:ARG:549: | B:SER:573: | 3.04 | A | B | Polar |
| A:ARG:550: | B:ASP:501: | 2.94 | A | B | Polar |
| A:ARG:550: | B:GLY:505: | 3.4 | A | B | Contact |
| A:ARG:550: | B:LEU:512: | 3.43 | A | B | Polar |
| A:ARG:550: | B:PRO:514: | 4.63 | A | B | Contact |
| A:ARG:550: | B:SER:510: | 3.54 | A | B | Polar |
| A:ARG:550: | B:TYR:511: | 2.81 | A | B | Contact |
| A:ARG:550: | B:TYR:513: | 3.37 | A | B | Contact |
| A:ARG:551: | B:TYR:511: | 3.06 | A | B | Contact |
| A:ASN:42: | B:ASN:219: | 3.23 | A | B | Polar |
| A:ASN:42: | B:LEU:117: | 3.67 | A | B | Contact |
| A:ASN:42: | B:LYS:120: | 4.24 | A | B | Contact |
| A:ASN:42: | B:PHE:116: | 3.45 | A | B | Contact |
| A:ASN:42: | B:TYR:113: | 3.34 | A | B | HBond,Polar |
| A:CYS:34: | B:GLN:218: | 4.49 | A | B | Contact |
| A:GLU:32: | B:LEU:249: | 3.17 | A | B | Contact |
| A:GLU:32: | B:LYS:328: | 2.62 | A | B | Polar |
| A:GLU:41: | B:LYS:120: | 2.72 | A | B | Polar |

| | | | | | |
|---|---|---|---|---|---|
| A:GLU:41: | B:PHE:116: | 4.3 | A | B | Contact |
| A:GLU:536: | B:LEU:512: | 3.83 | A | B | Contact |
| A:GLY:543: | B:SER:561: | 3.72 | A | B | Contact |
| A:GLY:543: | B:TYR:513: | 4.92 | A | B | Contact |
| A:GLY:543: | B:TYR:558: | 3.99 | A | B | Contact |
| A:ILE:39: | B:ASN:219: | 3.8 | A | B | Contact |
| A:ILE:39: | B:GLN:218: | 4.53 | A | B | Contact |
| A:ILE:39: | B:GLY:220: | 3.61 | A | B | Contact |
| A:ILE:542: | B:ASP:557: | 3.03 | A | B | Contact |
| A:ILE:542: | B:SER:561: | 4.35 | A | B | Contact |
| A:ILE:542: | B:VAL:567: | 4.43 | A | B | Contact |
| A:LEU:43: | B:ASP:382: | 4.79 | A | B | Contact |
| A:LEU:43: | B:TYR:113: | 4.46 | A | B | Contact |
| A:LEU:49: | B:GLN:109: | 3.52 | A | B | Contact |
| A:LEU:49: | B:LEU:112: | 4.55 | A | B | Contact |
| A:LEU:49: | B:TYR:113: | 3.62 | A | B | Contact |
| A:LEU:51: | B:GLN:109: | 3.44 | A | B | Contact |
| A:LEU:51: | B:ILE:383: | 3.68 | A | B | Contact |
| A:LEU:51: | B:LEU:106: | 3.83 | A | B | Contact |
| A:LEU:51: | B:THR:110: | 3.42 | A | B | Contact |
| A:LEU:51: | B:TYR:113: | 3.6 | A | B | Contact |
| A:LEU:52: | B:ASP:382: | 3.44 | A | B | Contact |
| A:LEU:52: | B:ILE:383: | 4.79 | A | B | Contact |
| A:LEU:546: | B:ARG:569: | 3.45 | A | B | Contact |
| A:LEU:546: | B:ASP:554: | 3.4 | A | B | Contact |
| A:LEU:546: | B:ASP:557: | 3.48 | A | B | Contact |
| A:LEU:546: | B:TYR:513: | 4.07 | A | B | Contact |
| A:LEU:546: | B:TYR:558: | 3.6 | A | B | Contact |
| A:LEU:54: | B:GLN:109: | 3.68 | A | B | Contact |
| A:LEU:54: | B:GLU:105: | 3.54 | A | B | Contact |
| A:LEU:54: | B:LEU:106: | 3.7 | A | B | Contact |
| A:LEU:58: | B:GLN:99: | 3.89 | A | B | Contact |
| A:LEU:58: | B:LEU:103: | 3.67 | A | B | Contact |
| A:LEU:58: | B:LEU:106: | 3.18 | A | B | Contact |
| A:LEU:58: | B:LYS:102: | 3.74 | A | B | Contact |
| A:LEU:58: | B:TRP:393: | 3.23 | A | B | Polar |
| A:LYS:57: | B:LYS:102: | 4.04 | A | B | Contact |
| A:LYS:59: | B:GLU:389: | 3.34 | A | B | Contact |
| A:LYS:59: | B:THR:385: | 2.92 | A | B | Polar |
| A:LYS:59: | B:TRP:393: | 4.23 | A | B | Contact |

| | | | | | |
|---|---|---|---|---|---|
| A:MET:46: | B:ASP:382: | 4.54 | A | B | Contact |
| A:MET:46: | B:ILE:383: | 3.85 | A | B | Contact |
| A:MET:46: | B:TYR:113: | 3.74 | A | B | Contact |
| A:PRO:537: | B:GLU:562: | 3.58 | A | B | Contact |
| A:PRO:537: | B:TYR:558: | 4.1 | A | B | Contact |
| A:PRO:60: | B:GLU:389: | 4.35 | A | B | Contact |
| A:PRO:60: | B:TRP:393: | 4.7 | A | B | Contact |
| A:SER:29: | B:ASN:247: | 4.11 | A | B | Contact |
| A:SER:541: | B:SER:561: | 4.52 | A | B | Contact |
| A:SER:55: | B:ASP:382: | 4.52 | A | B | Contact |
| A:SER:55: | B:ILE:383: | 2.67 | A | B | HBond,Polar |
| A:SER:55: | B:LEU:106: | 3.87 | A | B | Contact |
| A:SER:55: | B:PHE:384: | 4.88 | A | B | Contact |
| A:SER:55: | B:THR:385: | 4.47 | A | B | Contact |
| A:TYR:31: | B:ASN:247: | 3.97 | A | B | Contact |
| A:TYR:31: | B:GLN:218: | 2.85 | A | B | Polar |
| A:TYR:31: | B:HIS:243: | 2.52 | A | B | Polar |
| A:TYR:31: | B:ILE:129: | 3.51 | A | B | Contact |
| A:TYR:31: | B:ILE:214: | 3.78 | A | B | Contact |
| A:TYR:31: | B:LEU:244: | 4.74 | A | B | Contact |
| A:TYR:31: | B:LEU:249: | 4.2 | A | B | Contact |
| A:TYR:31: | B:TRP:217: | 3.16 | A | B | Contact |
| B:ARG:569: | A:ARG:549: | 4.96 | B | A | Contact |
| B:ARG:569: | A:LEU:546: | 3.45 | B | A | Contact |
| B:ASN:219: | A:ARG:38: | 4.32 | B | A | Contact |
| B:ASN:219: | A:ASN:42: | 3.23 | B | A | Polar |
| B:ASN:219: | A:ILE:39: | 3.8 | B | A | Contact |
| B:ASN:247: | A:SER:29: | 4.11 | B | A | Contact |
| B:ASN:247: | A:TYR:31: | 3.97 | B | A | Contact |
| B:ASN:356: | A:ARG:35: | 2.42 | B | A | Polar |
| B:ASP:355: | A:ARG:35: | 4.02 | B | A | Contact |
| B:ASP:382: | A:LEU:43: | 4.79 | B | A | Contact |
| B:ASP:382: | A:LEU:52: | 3.44 | B | A | Contact |
| B:ASP:382: | A:MET:46: | 4.54 | B | A | Contact |
| B:ASP:382: | A:SER:55: | 4.52 | B | A | Contact |
| B:ASP:501: | A:ARG:550: | 2.94 | B | A | Polar |
| B:ASP:554: | A:ARG:549: | 4.9 | B | A | Contact |
| B:ASP:554: | A:LEU:546: | 3.4 | B | A | Contact |
| B:ASP:557: | A:ILE:542: | 3.03 | B | A | Contact |
| B:ASP:557: | A:LEU:546: | 3.48 | B | A | Contact |

| | | | | | |
|---|---|---|---|---|---|
| B:ASP:571: | A:ARG:549: | 4.13 | B | A | Contact |
| B:GLN:109: | A:LEU:49: | 3.52 | B | A | Contact |
| B:GLN:109: | A:LEU:51: | 3.44 | B | A | Contact |
| B:GLN:109: | A:LEU:54: | 3.68 | B | A | Contact |
| B:GLN:218: | A:ARG:35: | 3.52 | B | A | Contact |
| B:GLN:218: | A:ARG:38: | 2.84 | B | A | Polar |
| B:GLN:218: | A:CYS:34: | 4.49 | B | A | Contact |
| B:GLN:218: | A:ILE:39: | 4.53 | B | A | Contact |
| B:GLN:218: | A:TYR:31: | 2.85 | B | A | Polar |
| B:GLN:99: | A:LEU:58: | 3.89 | B | A | Contact |
| B:GLU:105: | A:LEU:54: | 3.54 | B | A | Contact |
| B:GLU:132: | A:ARG:38: | 2.76 | B | A | Polar |
| B:GLU:389: | A:LYS:59: | 3.34 | B | A | Contact |
| B:GLU:389: | A:PRO:60: | 4.35 | B | A | Contact |
| B:GLU:562: | A:PRO:537: | 3.58 | B | A | Contact |
| B:GLY:128: | A:ARG:38: | 4.95 | B | A | Contact |
| B:GLY:220: | A:ARG:35: | 4.2 | B | A | Contact |
| B:GLY:220: | A:ILE:39: | 3.61 | B | A | Contact |
| B:GLY:505: | A:ARG:550: | 3.4 | B | A | Contact |
| B:HIS:243: | A:TYR:31: | 2.52 | B | A | Polar |
| B:ILE:124: | A:ARG:38: | 3.84 | B | A | Contact |
| B:ILE:129: | A:ARG:38: | 3.34 | B | A | Polar |
| B:ILE:129: | A:TYR:31: | 3.51 | B | A | Contact |
| B:ILE:214: | A:TYR:31: | 3.78 | B | A | Contact |
| B:ILE:383: | A:LEU:51: | 3.68 | B | A | Contact |
| B:ILE:383: | A:LEU:52: | 4.79 | B | A | Contact |
| B:ILE:383: | A:MET:46: | 3.85 | B | A | Contact |
| B:ILE:383: | A:SER:55: | 2.67 | B | A | HBond,Polar |
| B:LEU:103: | A:LEU:58: | 3.67 | B | A | Contact |
| B:LEU:106: | A:LEU:51: | 3.83 | B | A | Contact |
| B:LEU:106: | A:LEU:54: | 3.7 | B | A | Contact |
| B:LEU:106: | A:LEU:58: | 3.18 | B | A | Contact |
| B:LEU:106: | A:SER:55: | 3.87 | B | A | Contact |
| B:LEU:112: | A:LEU:49: | 4.55 | B | A | Contact |
| B:LEU:117: | A:ASN:42: | 3.67 | B | A | Contact |
| B:LEU:244: | A:TYR:31: | 4.74 | B | A | Contact |
| B:LEU:249: | A:GLU:32: | 3.17 | B | A | Contact |
| B:LEU:249: | A:TYR:31: | 4.2 | B | A | Contact |
| B:LEU:512: | A:ARG:547: | 3.27 | B | A | Contact |
| B:LEU:512: | A:ARG:550: | 3.43 | B | A | Polar |

| | | | | | |
|---|---|---|---|---|---|
| B:LEU:512: | A:GLU:536: | 3.83 | B | A | Contact |
| B:LYS:102: | A:LEU:58: | 3.74 | B | A | Contact |
| B:LYS:102: | A:LYS:57: | 4.04 | B | A | Contact |
| B:LYS:120: | A:ASN:42: | 4.24 | B | A | Contact |
| B:LYS:120: | A:GLU:41: | 2.72 | B | A | Polar |
| B:LYS:328: | A:GLU:32: | 2.62 | B | A | Polar |
| B:PHE:116: | A:ARG:45: | 3.61 | B | A | Contact |
| B:PHE:116: | A:ASN:42: | 3.45 | B | A | Contact |
| B:PHE:116: | A:GLU:41: | 4.3 | B | A | Contact |
| B:PHE:384: | A:SER:55: | 4.88 | B | A | Contact |
| B:PRO:514: | A:ARG:550: | 4.63 | B | A | Contact |
| B:SER:510: | A:ARG:550: | 3.54 | B | A | Polar |
| B:SER:561: | A:GLY:543: | 3.72 | B | A | Contact |
| B:SER:561: | A:ILE:542: | 4.35 | B | A | Contact |
| B:SER:561: | A:SER:541: | 4.52 | B | A | Contact |
| B:SER:573: | A:ARG:549: | 3.04 | B | A | Polar |
| B:THR:110: | A:LEU:51: | 3.42 | B | A | Contact |
| B:THR:385: | A:LYS:59: | 2.92 | B | A | Polar |
| B:THR:385: | A:SER:55: | 4.47 | B | A | Contact |
| B:TRP:217: | A:ARG:35: | 2.97 | B | A | HBond,Polar |
| B:TRP:217: | A:TYR:31: | 3.16 | B | A | Contact |
| B:TRP:393: | A:LEU:58: | 3.23 | B | A | Polar |
| B:TRP:393: | A:LYS:59: | 4.23 | B | A | Contact |
| B:TRP:393: | A:PRO:60: | 4.7 | B | A | Contact |
| B:TYR:113: | A:ARG:45: | 4.42 | B | A | Contact |
| B:TYR:113: | A:ASN:42: | 3.34 | B | A | HBond,Polar |
| B:TYR:113: | A:LEU:43: | 4.46 | B | A | Contact |
| B:TYR:113: | A:LEU:49: | 3.62 | B | A | Contact |
| B:TYR:113: | A:LEU:51: | 3.6 | B | A | Contact |
| B:TYR:113: | A:MET:46: | 3.74 | B | A | Contact |
| B:TYR:511: | A:ARG:547: | 3.54 | B | A | Contact |
| B:TYR:511: | A:ARG:550: | 2.81 | B | A | Contact |
| B:TYR:511: | A:ARG:551: | 3.06 | B | A | Contact |
| B:TYR:513: | A:ARG:547: | 3.97 | B | A | Contact |
| B:TYR:513: | A:ARG:550: | 3.37 | B | A | Contact |
| B:TYR:513: | A:GLY:543: | 4.92 | B | A | Contact |
| B:TYR:513: | A:LEU:546: | 4.07 | B | A | Contact |
| B:TYR:558: | A:ARG:547: | 4.31 | B | A | Contact |
| B:TYR:558: | A:GLY:543: | 3.99 | B | A | Contact |
| B:TYR:558: | A:LEU:546: | 3.6 | B | A | Contact |

| | | | | | |
|---|---|---|---|---|---|
| B:TYR:558: | A:PRO:537: | 4.1 | B | A | Contact |
| B:VAL:567: | A:ILE:542: | 4.43 | B | A | Contact |

**Supplementary Table 2: CDCA7B DDM1 interaction Residues**

| residue_1 | residue_2 | distance | chain_1 | chain_2 | interaction |
|---|---|---|---|---|---|
| A:ARG:35: | B:ASN:356: | 2.8 | A | B | Polar |
| A:ARG:35: | B:ASP:355: | 3.94 | A | B | Contact |
| A:ARG:35: | B:GLN:218: | 3.53 | A | B | Contact |
| A:ARG:35: | B:GLY:220: | 4.24 | A | B | Contact |
| A:ARG:35: | B:TRP:217: | 3.03 | A | B | HBond,Polar |
| A:ARG:38: | B:ASN:219: | 4.19 | A | B | Contact |
| A:ARG:38: | B:GLN:218: | 3.56 | A | B | Polar |
| A:ARG:38: | B:ILE:124: | 3.76 | A | B | Contact |
| A:ARG:38: | B:ILE:129: | 4.47 | A | B | Contact |
| A:ARG:394: | B:ASN:540: | 4.43 | A | B | Contact |
| A:ARG:394: | B:GLU:589: | 4.01 | A | B | Contact |
| A:ARG:394: | B:LYS:542: | 4.51 | A | B | Contact |
| A:ARG:394: | B:SER:591: | 4.13 | A | B | Contact |
| A:ARG:394: | B:SER:592: | 4.89 | A | B | Contact |
| A:ARG:394: | B:SER:594: | 4.29 | A | B | Contact |
| A:ARG:410: | B:GLU:562: | 2.6 | A | B | Polar |
| A:ARG:410: | B:LYS:563: | 4.84 | A | B | Contact |
| A:ARG:410: | B:SER:561: | 3.02 | A | B | Polar |
| A:ARG:412: | B:TYR:511: | 4.52 | A | B | Contact |
| A:ARG:413: | B:LEU:512: | 3.06 | A | B | Contact |
| A:ARG:413: | B:PRO:514: | 4.94 | A | B | Contact |
| A:ARG:413: | B:TYR:513: | 3.06 | A | B | Polar |
| A:ARG:413: | B:TYR:558: | 2.92 | A | B | Polar |
| A:ARG:45: | B:PHE:116: | 3.58 | A | B | Contact |
| A:ARG:45: | B:TYR:113: | 4.34 | A | B | Contact |
| A:ASN:42: | B:ASN:219: | 3.21 | A | B | Polar |
| A:ASN:42: | B:LEU:117: | 3.57 | A | B | Contact |
| A:ASN:42: | B:LYS:120: | 4.24 | A | B | Contact |
| A:ASN:42: | B:PHE:116: | 3.42 | A | B | Contact |
| A:ASN:42: | B:TYR:113: | 3.35 | A | B | HBond,Polar |
| A:CYS:34: | B:GLN:218: | 4.9 | A | B | Contact |
| A:GLN:416: | B:LEU:512: | 4.63 | A | B | Contact |
| A:GLU:32: | B:LEU:249: | 2.51 | A | B | Contact |
| A:GLU:32: | B:LYS:328: | 2.64 | A | B | Polar |
| A:GLU:32: | B:TRP:217: | 4.84 | A | B | Contact |
| A:GLU:41: | B:LYS:120: | 2.62 | A | B | Polar |
| A:GLU:41: | B:PHE:116: | 4.21 | A | B | Contact |
| A:GLU:61: | B:GLN:99: | 4.28 | A | B | Contact |

| | | | | | |
|---|---|---|---|---|---|
| A:GLU:61: | B:LYS:102: | 2.38 | A | B | Polar |
| A:GLY:406: | B:ASP:557: | 3.75 | A | B | Contact |
| A:GLY:406: | B:SER:561: | 3.9 | A | B | Contact |
| A:GLY:406: | B:TYR:558: | 4.26 | A | B | Contact |
| A:GLY:407: | B:SER:561: | 3.99 | A | B | Contact |
| A:ILE:39: | B:ASN:219: | 3.82 | A | B | Contact |
| A:ILE:39: | B:GLN:218: | 4.7 | A | B | Contact |
| A:ILE:39: | B:GLY:220: | 3.69 | A | B | Contact |
| A:ILE:405: | B:ARG:569: | 2.76 | A | B | Contact |
| A:ILE:405: | B:ASP:554: | 3.42 | A | B | Contact |
| A:ILE:405: | B:ASP:557: | 3.13 | A | B | HBond,Polar |
| A:ILE:51: | B:GLN:109: | 3.52 | A | B | Contact |
| A:ILE:51: | B:ILE:383: | 3.5 | A | B | Contact |
| A:ILE:51: | B:LEU:106: | 3.69 | A | B | Contact |
| A:ILE:51: | B:THR:110: | 3.89 | A | B | Contact |
| A:ILE:51: | B:TYR:113: | 3.49 | A | B | Contact |
| A:ILE:62: | B:GLN:99: | 4.75 | A | B | Contact |
| A:ILE:62: | B:GLU:389: | 4.42 | A | B | Contact |
| A:ILE:62: | B:SER:392: | 3.29 | A | B | Contact |
| A:ILE:62: | B:TRP:393: | 2.25 | A | B | Contact |
| A:LEU:251: | B:ALA:87: | 4.04 | A | B | Contact |
| A:LEU:409: | B:TYR:511: | 3.73 | A | B | Contact |
| A:LEU:409: | B:TYR:513: | 3.1 | A | B | Contact |
| A:LEU:409: | B:TYR:558: | 4.42 | A | B | Contact |
| A:LEU:43: | B:TYR:113: | 4.43 | A | B | Contact |
| A:LEU:49: | B:GLN:109: | 3.58 | A | B | Contact |
| A:LEU:49: | B:LEU:112: | 4.41 | A | B | Contact |
| A:LEU:49: | B:TYR:113: | 3.66 | A | B | Contact |
| A:LEU:54: | B:GLN:109: | 3.83 | A | B | Contact |
| A:LEU:54: | B:GLU:105: | 3.72 | A | B | Contact |
| A:LEU:54: | B:LEU:106: | 3.88 | A | B | Contact |
| A:LEU:58: | B:LEU:103: | 3.93 | A | B | Contact |
| A:LEU:58: | B:LEU:106: | 3.84 | A | B | Contact |
| A:LEU:58: | B:LYS:102: | 3.82 | A | B | Contact |
| A:LEU:58: | B:TRP:393: | 2.97 | A | B | Contact |
| A:LYS:247: | B:GLN:480: | 3.61 | A | B | Contact |
| A:LYS:395: | B:SER:592: | 4.26 | A | B | Contact |
| A:LYS:59: | B:ASP:382: | 4.21 | A | B | Contact |
| A:LYS:59: | B:GLU:389: | 3.21 | A | B | Contact |
| A:LYS:59: | B:ILE:383: | 4.11 | A | B | Contact |

| | | | | | |
|---|---|---|---|---|---|
| A:LYS:59: | B:THR:385: | 2.8 | A | B | Polar |
| A:MET:396: | B:GLY:564: | 3.71 | A | B | Contact |
| A:MET:396: | B:LYS:563: | 4.52 | A | B | Contact |
| A:MET:396: | B:PHE:565: | 3.51 | A | B | Contact |
| A:MET:46: | B:ASP:382: | 4.98 | A | B | Contact |
| A:MET:46: | B:ILE:383: | 3.84 | A | B | Contact |
| A:MET:46: | B:TYR:113: | 3.79 | A | B | Contact |
| A:MET:52: | B:ASP:382: | 4.13 | A | B | Contact |
| A:PRO:252: | B:ALA:87: | 4.84 | A | B | Contact |
| A:PRO:400: | B:SER:561: | 3.87 | A | B | Contact |
| A:SER:29: | B:ASN:247: | 4.24 | A | B | Contact |
| A:SER:397: | B:GLU:566: | 2.16 | A | B | Polar |
| A:SER:397: | B:GLY:564: | 4.33 | A | B | Contact |
| A:SER:404: | B:ASP:557: | 4.28 | A | B | Contact |
| A:SER:404: | B:SER:561: | 4.19 | A | B | Contact |
| A:SER:55: | B:ASP:382: | 4.37 | A | B | Contact |
| A:SER:55: | B:ILE:383: | 2.85 | A | B | HBond,Polar |
| A:SER:55: | B:LEU:106: | 4.01 | A | B | Contact |
| A:TYR:31: | B:ASN:247: | 3.98 | A | B | Contact |
| A:TYR:31: | B:GLN:218: | 3.11 | A | B | Polar |
| A:TYR:31: | B:HIS:243: | 2.69 | A | B | Polar |
| A:TYR:31: | B:ILE:129: | 3.32 | A | B | Contact |
| A:TYR:31: | B:ILE:214: | 3.67 | A | B | Contact |
| A:TYR:31: | B:LEU:244: | 4.83 | A | B | Contact |
| A:TYR:31: | B:LEU:249: | 4.25 | A | B | Contact |
| A:TYR:31: | B:TRP:217: | 3 | A | B | Contact |
| A:VAL:140: | B:VAL:91: | 4.04 | A | B | Contact |
| B:ALA:87: | A:LEU:251: | 4.04 | B | A | Contact |
| B:ALA:87: | A:PRO:252: | 4.84 | B | A | Contact |
| B:ARG:569: | A:ILE:405: | 2.76 | B | A | Contact |
| B:ASN:219: | A:ARG:38: | 4.19 | B | A | Contact |
| B:ASN:219: | A:ASN:42: | 3.21 | B | A | Polar |
| B:ASN:219: | A:ILE:39: | 3.82 | B | A | Contact |
| B:ASN:247: | A:SER:29: | 4.24 | B | A | Contact |
| B:ASN:247: | A:TYR:31: | 3.98 | B | A | Contact |
| B:ASN:356: | A:ARG:35: | 2.8 | B | A | Polar |
| B:ASN:540: | A:ARG:394: | 4.43 | B | A | Contact |
| B:ASP:355: | A:ARG:35: | 3.94 | B | A | Contact |
| B:ASP:382: | A:LYS:59: | 4.21 | B | A | Contact |
| B:ASP:382: | A:MET:46: | 4.98 | B | A | Contact |

| | | | | | |
|---|---|---|---|---|---|
| B:ASP:382: | A:MET:52: | 4.13 | B | A | Contact |
| B:ASP:382: | A:SER:55: | 4.37 | B | A | Contact |
| B:ASP:554: | A:ILE:405: | 3.42 | B | A | Contact |
| B:ASP:557: | A:GLY:406: | 3.75 | B | A | Contact |
| B:ASP:557: | A:ILE:405: | 3.13 | B | A | HBond,Polar |
| B:ASP:557: | A:SER:404: | 4.28 | B | A | Contact |
| B:GLN:109: | A:ILE:51: | 3.52 | B | A | Contact |
| B:GLN:109: | A:LEU:49: | 3.58 | B | A | Contact |
| B:GLN:109: | A:LEU:54: | 3.83 | B | A | Contact |
| B:GLN:218: | A:ARG:35: | 3.53 | B | A | Contact |
| B:GLN:218: | A:ARG:38: | 3.56 | B | A | Polar |
| B:GLN:218: | A:CYS:34: | 4.9 | B | A | Contact |
| B:GLN:218: | A:ILE:39: | 4.7 | B | A | Contact |
| B:GLN:218: | A:TYR:31: | 3.11 | B | A | Polar |
| B:GLN:480: | A:LYS:247: | 3.61 | B | A | Contact |
| B:GLN:99: | A:GLU:61: | 4.28 | B | A | Contact |
| B:GLN:99: | A:ILE:62: | 4.75 | B | A | Contact |
| B:GLU:105: | A:LEU:54: | 3.72 | B | A | Contact |
| B:GLU:389: | A:ILE:62: | 4.42 | B | A | Contact |
| B:GLU:389: | A:LYS:59: | 3.21 | B | A | Contact |
| B:GLU:562: | A:ARG:410: | 2.6 | B | A | Polar |
| B:GLU:566: | A:SER:397: | 2.16 | B | A | Polar |
| B:GLU:589: | A:ARG:394: | 4.01 | B | A | Contact |
| B:GLY:220: | A:ARG:35: | 4.24 | B | A | Contact |
| B:GLY:220: | A:ILE:39: | 3.69 | B | A | Contact |
| B:GLY:564: | A:MET:396: | 3.71 | B | A | Contact |
| B:GLY:564: | A:SER:397: | 4.33 | B | A | Contact |
| B:HIS:243: | A:TYR:31: | 2.69 | B | A | Polar |
| B:ILE:124: | A:ARG:38: | 3.76 | B | A | Contact |
| B:ILE:129: | A:ARG:38: | 4.47 | B | A | Contact |
| B:ILE:129: | A:TYR:31: | 3.32 | B | A | Contact |
| B:ILE:214: | A:TYR:31: | 3.67 | B | A | Contact |
| B:ILE:383: | A:ILE:51: | 3.5 | B | A | Contact |
| B:ILE:383: | A:LYS:59: | 4.11 | B | A | Contact |
| B:ILE:383: | A:MET:46: | 3.84 | B | A | Contact |
| B:ILE:383: | A:SER:55: | 2.85 | B | A | HBond,Polar |
| B:LEU:103: | A:LEU:58: | 3.93 | B | A | Contact |
| B:LEU:106: | A:ILE:51: | 3.69 | B | A | Contact |
| B:LEU:106: | A:LEU:54: | 3.88 | B | A | Contact |
| B:LEU:106: | A:LEU:58: | 3.84 | B | A | Contact |

| | | | | | |
|---|---|---|---|---|---|
| B:LEU:106: | A:SER:55: | 4.01 | B | A | Contact |
| B:LEU:112: | A:LEU:49: | 4.41 | B | A | Contact |
| B:LEU:117: | A:ASN:42: | 3.57 | B | A | Contact |
| B:LEU:244: | A:TYR:31: | 4.83 | B | A | Contact |
| B:LEU:249: | A:GLU:32: | 2.51 | B | A | Contact |
| B:LEU:249: | A:TYR:31: | 4.25 | B | A | Contact |
| B:LEU:512: | A:ARG:413: | 3.06 | B | A | Contact |
| B:LEU:512: | A:GLN:416: | 4.63 | B | A | Contact |
| B:LYS:102: | A:GLU:61: | 2.38 | B | A | Polar |
| B:LYS:102: | A:LEU:58: | 3.82 | B | A | Contact |
| B:LYS:120: | A:ASN:42: | 4.24 | B | A | Contact |
| B:LYS:120: | A:GLU:41: | 2.62 | B | A | Polar |
| B:LYS:328: | A:GLU:32: | 2.64 | B | A | Polar |
| B:LYS:542: | A:ARG:394: | 4.51 | B | A | Contact |
| B:LYS:563: | A:ARG:410: | 4.84 | B | A | Contact |
| B:LYS:563: | A:MET:396: | 4.52 | B | A | Contact |
| B:PHE:116: | A:ARG:45: | 3.58 | B | A | Contact |
| B:PHE:116: | A:ASN:42: | 3.42 | B | A | Contact |
| B:PHE:116: | A:GLU:41: | 4.21 | B | A | Contact |
| B:PHE:565: | A:MET:396: | 3.51 | B | A | Contact |
| B:PRO:514: | A:ARG:413: | 4.94 | B | A | Contact |
| B:SER:392: | A:ILE:62: | 3.29 | B | A | Contact |
| B:SER:561: | A:ARG:410: | 3.02 | B | A | Polar |
| B:SER:561: | A:GLY:406: | 3.9 | B | A | Contact |
| B:SER:561: | A:GLY:407: | 3.99 | B | A | Contact |
| B:SER:561: | A:PRO:400: | 3.87 | B | A | Contact |
| B:SER:561: | A:SER:404: | 4.19 | B | A | Contact |
| B:SER:591: | A:ARG:394: | 4.13 | B | A | Contact |
| B:SER:592: | A:ARG:394: | 4.89 | B | A | Contact |
| B:SER:592: | A:LYS:395: | 4.26 | B | A | Contact |
| B:SER:594: | A:ARG:394: | 4.29 | B | A | Contact |
| B:THR:110: | A:ILE:51: | 3.89 | B | A | Contact |
| B:THR:385: | A:LYS:59: | 2.8 | B | A | Polar |
| B:TRP:217: | A:ARG:35: | 3.03 | B | A | HBond,Polar |
| B:TRP:217: | A:GLU:32: | 4.84 | B | A | Contact |
| B:TRP:217: | A:TYR:31: | 3 | B | A | Contact |
| B:TRP:393: | A:ILE:62: | 2.25 | B | A | Contact |
| B:TRP:393: | A:LEU:58: | 2.97 | B | A | Contact |
| B:TYR:113: | A:ARG:45: | 4.34 | B | A | Contact |
| B:TYR:113: | A:ASN:42: | 3.35 | B | A | HBond,Polar |

| | | | | | |
|---|---|---|---|---|---|
| B:TYR:113: | A:ILE:51: | 3.49 | B | A | Contact |
| B:TYR:113: | A:LEU:43: | 4.43 | B | A | Contact |
| B:TYR:113: | A:LEU:49: | 3.66 | B | A | Contact |
| B:TYR:113: | A:MET:46: | 3.79 | B | A | Contact |
| B:TYR:511: | A:ARG:412: | 4.52 | B | A | Contact |
| B:TYR:511: | A:LEU:409: | 3.73 | B | A | Contact |
| B:TYR:513: | A:ARG:413: | 3.06 | B | A | Polar |
| B:TYR:513: | A:LEU:409: | 3.1 | B | A | Contact |
| B:TYR:558: | A:ARG:413: | 2.92 | B | A | Polar |
| B:TYR:558: | A:GLY:406: | 4.26 | B | A | Contact |
| B:TYR:558: | A:LEU:409: | 4.42 | B | A | Contact |
| B:VAL:91: | A:VAL:140: | 4.04 | B | A | Contact |
